# Dual blockade of misfolded alpha-sarcoglycan degradation by bortezomib and givinostat combination

**DOI:** 10.1101/2022.03.16.484594

**Authors:** Lucile Hoch, Nathalie Bourg, Fanny Degrugillier, Céline Bruge, Manon Benabides, Emilie Pellier, Johana Tournois, Gurvan Mahé, Nicolas Maignan, Jack Dawe, Maxime Georges, David Papazian, Nik Subramanian, Stéphanie Simon, Pascale Fanen, Cédric Delevoye, Isabelle Richard, Xavier Nissan

## Abstract

Limb-girdle muscular dystrophy type R3 (LGMD R3) is a rare genetic disorder characterized by a progressive proximal muscle weakness and caused by mutations in the *SGCA* gene encoding alpha-sarcoglycan (α-SG). Here, we report the results of a mechanistic screening ascertaining the molecular mechanisms involved in the degradation of the most prevalent misfolded R77C-α-SG protein. We performed a combinatorial study to identify drugs potentializing the effect of a low dose of the proteasome inhibitor bortezomib on the R77C-α-SG degradation inhibition. Analysis of the screening associated to artificial intelligence-based predictive ADMET characterization of the hits led to identification of the HDAC inhibitor givinostat as potential therapeutical candidate. Functional characterization revealed that givinostat effect was related to autophagic pathway inhibition, unveiling new theories concerning degradation pathways of misfolded SG proteins. Beyond the identification of a new therapeutic option for LGMD R3 patients, our results shed light on the potential repurposing of givinostat for the treatment of other genetic diseases sharing similar protein degradation defects such as LGMD R5 and cystic fibrosis.

## Introduction

Limb girdle muscular dystrophies (LGMDs) are a diverse group of genetic muscular dystrophies sharing common features of weakness and wasting of the shoulder and pelvic girdle muscles (Taghizadeh et al., 2019). More than 30 different subtypes of LGMDs have been described with diverse severities depending on the affected gene (Straub et al., 2018). Sarcoglycanopathies are a subgroup of LGMDs caused by autosomal recessive mutations in genes encoding the four sarcoglycan (SG) proteins, α–, β–, δ– and γ-SG, leading to LGMD R3 (Roberds et al., 1994), LGMD R4 (Bönnemann et al., 1995; Lim et al., 1995), LGMD R5 (Noguchi et al., 1995) and LGMD R6 (Nigro et al., 1996), respectively. SGs form a transmembrane complex playing a key role to protect striated muscles membrane against contraction-induced damages.

In the past two decades, more than 170 different mutations were described (Sandonà and Betto, 2009) to cause sarcoglycanopathies, including 70 *SGCA* gene mutations leading to LGMD R3. The most frequent, the substitution R77C (Carrié et al., 1997), leads to the expression of a misfolded α-SG protein which is recognized by the endoplasmic reticulum quality control (ERQC) system and subsequently degraded by the endoplasmic reticulum associated degradation (ERAD) system involving the proteasome (Bartoli et al., 2008; Gastaldello et al., 2008). Even if there is still no treatment available for LGMDs, preclinical investigations have recently described the positive impact of gene therapy approaches using AAV-mediated *SGCA* or *SGCG* gene transfer to rescue α-SG or γ-SG deficiencies (Mendell et al., 2009, 2010; Israeli et al., 2019). Recently, the results of a phase 1/2 clinical trial enrolling six LGMD R3 patients were reported showing a good tolerance and delivery efficacy of AAVrh74 in muscles but limited functional outcome improvements in patients (Mendell et al., 2019).

Several pharmacological approaches were also investigated to restore α-SG expression and function at the plasma membrane (Bartoli et al., 2008; Gastaldello et al., 2008; Soheili et al., 2012; Carotti et al., 2018). Most of these approaches were based on mechanistic hypothesis focused on missense mutations, especially on the two most frequently reported missense α-SG mutations R77C and V247M (Bartoli et al., 2008; Gastaldello et al., 2008). Even if mutated, the corresponding proteins were shown to be functional at the membrane level, opening the possibility of recovery strategies. To date, the main therapeutic avenues for these mutations are to target either the degradative proteins pathway comprising ERQC system and proteasome or the protein folding/maturation pathway with small molecules as revealed by the use of MG132 (Gastaldello et al., 2008), kifunensine (Soheili et al., 2012) or cystic fibrosis transmembrane regulator (CFTR) correctors (Carotti et al., 2018). Targeting the proteasomal pathway has pioneerly been developed for the treatment of multiple myeloma and lymphoma to induce cancer cells death and has led to the development of the FDA-approved proteasome inhibitors, carfilzomib (CFZ) or bortezomib (BTZ). However, while being potent proteasome inhibitors, these treatments are associated to side effect as peripheral neuropathy, thrombocytopenia, leukopenia, gastrointestinal dysfunction and hematological toxicity precluding their long-term use for muscular dystrophies (Kaplan et al., 2017; Mohan et al., 2017).

Recent developments in rare diseases highlighted the interest to perform drug screening to repurpose new drugs as revealed with the discovery of small molecules modulators of SMN expression for spinal muscular atrophy (SMA) (Jarecki et al., 2005), cardiac glycosides as modulators of myotonic dystrophy 1 (DM1) (Maury et al., 2019) or metformin as a modulator of progerin expression for Hutchinson–Gilford progeria syndrome (HGPS) (Egesipe et al., 2016). Recently, our group developed a pharmacological model of LGMD R3 to identify drugs rescuing the proteasomal degradation of R77C-α-SG. We previously reported that among 2000 tested drugs only one drug, thiostrepton, was capable to rescue R77C-α-SG membrane localization via a direct inhibition of the proteasome (Hoch et al., 2019). Here we have extended the library to 958 other drugs and performed the screening of these drugs alone or in combination with a low dose of BTZ. Our results demonstrate that the combination of BTZ with several heat shock protein 90 (HSP90) and histone deacetylase (HDAC) inhibitors exhibited potent capacity to restore the expression of the α-SG mutant form in the plasma membrane. The functional characterization of the HDAC inhibitor givinostat, revealed the inhibition of the autophagic degradation opening new strategies of combined treatments for LGMD R3.

## Materials and Methods

### Cell Culture

The R77C fibroblasts used in this study were isolated from a patient biopsy obtained by the Genethon’s Cell Bank. Informed consents were obtained from the parents of the patient included in this study, complying with the ethical guidelines of the institutions involved and with the legislation requirements of the country of origin. The R77C fibroblasts were transduced with a pBABE-puro-based retroviral vector containing sequence encoding the catalytic subunit of human telomerase reverse transcriptase (hTERT) and then selected in the presence of puromycin (1 mg/ml) for 10 days, as previously described (Hoch et al., 2019). Experimental protocols were approved by the french minister of research: AC-2013-1868. The SGCG KO fibroblasts used in this study were isolated from a biopsy obtained by the Genethon’s Cell Bank from a patient carrying the mutation c.525del at the homozygous state. Both fibroblast lines were cultured in Dulbecco’s modified Eagle’s medium+GlutaMAX (Invitrogen) supplemented with 10% fetal bovine serum (research grade, Sigma-Aldrich) and 1% Penicillin-Streptomycin (Invitrogen). For overexpression of the mutated α-SG or mutated γ-SG, fibroblasts were transduced with a lentiviral vector expressing human R77C-, R34H-, I124T-, V247M-α-SGmCh or C283Y-, E263K-γ-SGmCh with a multiplicity of infection (MOI) of 20 in the presence of 4 μg/ml of polybrene (Sigma-Aldrich). Cells were seeded on plates coated with 50μg/ml of collagen I and maintained in a humidified atmosphere of 5% CO_2_ at 37° C. Human embryonic kidney (HEK293 MSR Grip Tite) cells stably coexpressing eYFP (H148Q/I152L) and F508del-CFTR-3HA (HEK-F508del/YFP) were grown in DMEM supplemented with 10% FBS, 50 μg/mL zeocin, 0.6 mg/mL G418, 10 μg/mL blasticidin. Human bronchial epithelial cell line CFBE41o-derived from a CF patient were modified to stably expressing F508del CFTR (F508del-CFTR) (Kunzelmann et al., 1993; Gruenert et al., 2004; Illek et al., 2008). CFBE cells were grown in MEM (Life Technologies) supplemented with 10% FBS (Sigma Aldrich), 1% penicillin/streptomycin (P/S) (Invitrogen) and 1% L-glutamine (Invitrogen).

### Plasmid cloning and mutagenesis

The sarcoglycan fusion protein was designed based on the human *SGCA* and *SGCG* consensus coding sequence (CCDS) found in the NCBI portal (Gene ID: 6442, CCDS number 45729.1 and Gene ID: 6445, CCDS number 9299.1) and the mCherry sequence. The linker, in the amino-acid form of -GGGGS-, was chosen as a flexible type linker that also increased stability/folding of the fusion proteins (Hoch et al., 2019). The fusion protein nucleotide sequence was synthesized by Genecust and cloned in a vector plasmid prepared for lentivirus production. The final construct, termed α-SGmCh or γ-SGmCh, were driven by the cytomegalovirus (CMV) promoter. The R34H, R77C, I124T and V247M mutations were generated based on the 100GT > AC, 229C > T, 371T > C and 739G > A nucleotide changes of *SGCA* gene, respectively. The E263K and C283Y mutations were generated based on the 788G > A and 847C > A nucleotide changes of *SGCG* gene, respectively. Mutagenesis was performed using the QuikChange XL Site-Directed Mutagenesis kit (200516, Agilent) according to manufacturer’s instructions. The primers used to introduce the R34H, R77C, I124T, V247M, E263K and C283Y mutations are listed in Supplementary Table 1.

### Pharmacological treatments

Twenty-four hours after seeding, cells were treated with BTZ (Selleckchem), CFZ (Selleckchem), MG132 (Selleckchem), givinostat (Selleckchem), belinostat (Selleckchem), tubacin (Selleckchem), Lumacaftor (VX-809; Selleckchem), Ivacaftor (VX-770; Selleckchem) or the carrier 0.1% dimethyl sulfoxide (DMSO, VWR). Cells were analyzed after 24 hours of treatment.

### High throughput screening

The high-throughput screening was conducted on Biocell 1800 (Agilent). For this, 1600 R77C-α-SGmCh fibroblasts were seeded in 36 μl of culture medium into black 384-well clear bottom plates. Twenty-four hours after seeding, 2 μl of compounds from the chemical library were transferred in monoplicate into cell assay plates added with 2 μl of culture medium for non-combined treatments or 5 nM BTZ for combined treatments. In each plate, negative control (0.1% DMSO) and positive control (30 nM BTZ) were added in columns 1-23 and upper half columns 2-24, respectively. 5 nM BTZ was also added in the lower half columns 2-24 to control the low effect of this concentration during the drug screening. Plates were then incubated for 24 hours and processed for α-SG detection assay. Each of the 6 plates of the screening were fixed and stained with the specific anti-α-SG antibody using the optimized protocol under non-permeabilized condition. To prevent the discovery of toxic molecules, the number of cells was monitored in parallel by counting Hoechst stained cells per field and candidates showing mortality superior to 55 % were excluded.

### SG localization cell-based assay

After 24 hours of drug treatment, cells were fixed in 4% paraformaldehyde (10 min, room temperature). Immunocytochemistry was performed in a phosphate-buffered saline (PBS) solution supplemented with 1% bovine serum albumin (BSA; Sigma) for blocking (1 hour, room temperature) and with a mouse anti-α-SG 1:100 (NCL-L-a-SARC, Novocastra, Leica) or a mouse anti-γ-SG 1:100 (NCL-L-g-SARC, Novocastra, Leica) for primary hybridization step (overnight, 4 °C). Cells were stained with a fluorophore-conjugated secondary anti-mouse antibody (Invitrogen; 1 hour, room temperature) and nuclei were visualized with Hoechst 33342 (Invitrogen). SG localization was analyzed with a CellInsight CX7 HCS Platform (Cellomics Inc). The first channel was used for nuclei identification, the second one for membrane α-SG or γ-SG staining identification and the third one for mCherry tag identification. Pictures were acquired with a 10x objective in high-resolution camera mode and were analyzed using the colocalization bioapplication quantifying positive cells for membrane α-SG or γ-SG staining and mCherry tag expression.

### Chemical library

The chemical library was obtained from Selleckchem, distributed in 384 well plate format and tested at 5 μM. The Selleckchem Chemical library contains 958 FDA approved drugs and pharmacologically active compounds.

### Data analysis

Data analysis of the screening was performed using a customized Hiscreen application (Discngine) connected to Spotfire software (Tibco Software Inc.). The robustness of the assay was evaluated by calculating for each plate the *Z*Δ factor on the percentage of mCherry/α-SG positive cells parameter as follows Z□=1–[3(SDP+SDN)/(MP– MN)] where MP and MN correspond to the means of the positive (30 nM BTZ) and negative (0.1% DMSO) controls, respectively, and SDP and SDN correspond to their S.D. Raw data related to the percentage of mCherry/α-SG positive cells were normalized to the mean of positive and negative controls, which are defined as 100% and 0%, respectively. Raw data related to cell number per field were normalized to the mean of negative controls. Hit selection was performed using in parallel the number of standard deviations from the mean for each readout value (Z-score) calculated per plate individually or per run where all plate data were pooled. Only hits whose Z-score plate and/or Z-score run was ⍰2 and that did not decrease cell number by more than 55% compared to 0.1% DMSO condition were selected for subsequent validation step. Hits were then tested at gradual concentrations (10 nM - 10 μM) for parallel exploration of their efficacy and toxicity.

### ADMET analysis

76 ADMET endpoints were evaluated for all the compounds found to be active on the assay. These 76 properties represent a set of drug-likeness parameters that are used to compare compounds. Among these parameters, 27 can be directly determined from the sole structure of the considered molecules. The remaining 49 parameters are normally evaluated through assays. Therefore, assay data was collected for approximately 150k compounds and Machine Learning models were trained to predict assay results using molecules features and descriptors and were evaluated through K-Fold train-test-splits. The predictions were then passed through so-called druglikeness decision rules, all determined from literature and the analysis of the properties of authorized drugs. Finally, a metric was created to establish which compounds maximize the number of passing ADMET endpoints, the viability of the cellular culture during the assay and the activity observed on the biomarker of the assay. The final ranking established from this metric is to be considered as a risk-benefit indicator highlighting the compounds being the most likely to both show efficacy and minimize adverse effects.

### Quantitative PCR

Total RNAs were isolated using the RNeasy Mini extraction kit (Qiagen) according to the manufacturer’s protocol. A DNase I digestion was performed to avoid genomic DNA amplification. RNA levels and quality were checked using the NanoDrop technology. A total of 500 ng of RNA was used for reverse transcription using the SuperScript III reverse transcription kit (Invitrogen). Quantitative polymerase chain reaction (qPCR) analysis was performed using a QuantStudio 12 K Flex real-time PCR system (Applied biosystem) and Luminaris Probe qPCR Master Mix (Thermo Scientific), following the manufacturers’ instructions. α-SG expression analysis were performed using the TaqMan gene expression Master Mix (Roche), following the manufacturer’s protocol. Quantification of gene expression was based on the DeltaCt method and normalized on 18S expression (Assay HS_099999). Primer sequences for endogenous *SGCA* were: 5’-TAAACAAGCAGGGAGAGGGG-3’ and 5’-CAATTGGTGAGCAGAGCAGC-3’. Primer sequences for exogenous *SGCA* were: 5’-TGCTGGCCTATGTCATGTGC-3’ and 5’-TCTGGATGTCGGAGGTAGCC-3’. Taqman MGB probe sequence for exogenous *SGCA* was 5’-CGGGAGGGAAGGCTGAAGAGAGACC-3’.

### Immunoprecipitation

Immunoprecipitations were performed using Dynabeads Protein A (Invitrogen) following the manufacturer’s protocol. Briefly, the HSP90 antibody (4877S, Cell Signaling) was preincubated with the beads for 30 minutes. The bead/antibody mix was then added to 100 μg of whole-cell lysate and incubated for additional 30 minutes. Samples were washed three times with PBS containing 0,1% Tween-20 and subjected to Western immunoblotting analysis.

### Western immunoblotting

Twenty-four hours after treatments with the tested compounds or the carrier, the immortalized fibroblasts expressing R77C-α-SGmCh or the CFBE-F508del-CFTR were collected. Proteins were extracted by cell lysis buffer (NP40 or RIPA Buffer, respectively, Thermo Scientific) and completed with Proteases Inhibitors (Complete PIC, Roche). Protein concentration was measured using the Pierce BCA Protein Assay Kit (ThermoScientific) and the absorbance at 562 nm was evaluated using a CLARIOstar^®^ microplate reader (BMG Labtech) or a TriStar plate reader (Berthold). For the fibroblasts extracts, a total of 20 μg of protein was separated using a 3–8% Criterion™ XT tris-acetate or a 4–15% Criterion™ XT tris-glycine protein gels and then transferred to PVDF membrane with a Trans-Blot Turbo Transfert system (Biorad) following the manufacturer’s instructions. For the CFBE extracts, a total of 50 μg of protein was separated on 7% sodium dodecyl sulfate polyacrylamide gel electrophoresis and then transferred onto nitrocellulose membrane (GE Healthcare) with a Novex Gel Transfert Device (ThermoScientific) following the manufacturer’s instructions. Membranes were then blocked in Odyssey blocking buffer (Li-Cor) or Tris-Buffer-Saline 1X (TBS) containing 0.05% Tween-20 and 5% milk for 1 hour at room temperature. Incubation with primary antibodies diluted in blocking buffer was carried out at 4°C from 2 hours to overnight for the mouse anti-α-SG 1:100 (NCL-L-a-SARC, Novocastra, Leica), the rabbit anti-HSP90 1:1000 (4877S, Cell Signaling), the rabbit anti-acetylated-lysine 1:1000 (9441S, Cell Signaling), the mouse anti-ubiquitinated proteins 1:1000 (04-263, Merck), the rabbit anti-LC3B 1:1000 (NB600-1384, Novus), the mouse anti-P62 1:1000 (ab56416, Abcam), the mouse anti-acetylated-α-tubulin 1:1000 (T6793, Sigma), the rabbit anti-V-ATPase0a1 (V0a1) 1:500 (NBP1-89342, Novus), the mouse anti-CFTR 1:5000 and 1:1000 (596 and 570, Cystic Fibrosis Foundation distribution program), the rabbit anti-LaminB1 1:15000 (ab16048, abcam) and for the rabbit anti-β-actin 1:1000 (926-42210, Li-Cor). Washing was carried out 4 times for 8 minutes at room temperature with TBS + 0.1% Tween 20 and the membranes were incubated with a donkey antimouse antibody IRDye-680 1:10000 (926-32222, Li-Cor) or a donkey anti-rabbit antibody IRDye-800 1:5000 (926-32213, Li-Cor) in blocking buffer. Washing was carried out 4 times for 8 minutes at room temperature with TBS + 0.1% Tween20. Proteins were detected by fluorescence (Odyssey, Li-Cor) or revealed using the ECL Prime Western Blotting Detection Reagent kit (GE Healthcare) and images were acquired in a dark room using a G:Box (Syngene) equipped with GeneSys software following the manufacturer’s instructions.

### Proteasomal activity assay

Fibroblasts were seeded in 384 well plates and treated with various concentrations of BTZ (100 pM - 300 nM), givinostat (10 nM - 10 μM), belinostat (10 nM - 10 μM), MG132 (30 nM - 10 μM) or CFZ (100 pM - 1 μM) for 24 hours. Proteasome-Glo™ chymotrypsin-like cell-based assay reagents were added according to manufacturer instructions (Promega). Luminescence was read using a CLARIOstar® microplate reader (BMG Labtech).

### Immunostaining assay

R77C-α-SGmCh fibroblasts were seeded on coverslips coated with with 50μg/ml of collagen I. After 24 hours, cells were treated with compounds during 24 hours and fixed in 4% formaldehyde. Permeabilization was performed with 0.5% Triton X-100 (ThermoScientific) for 5 min. Then, PBS containing 1% BSA was used for blocking. Incubation with primary antibodies diluted in blocking buffer was carried out at 4°C overnight for the mouse anti-KDEL 1:100 (ADI-SPA-827, Enzolife) and the mouse anti-LAMP2 1:500 (ab25631, Abcam). Washing was carried out 3 times for 5 minutes at room temperature with PBS and then secondary antibodies alexa fluor 488 goat anti-mouse 1:1000 (A-11029, ThermoScientific), alexa fluor 555 goat anti-rabbit 1:1000 (A-21429, ThermoScientific) and Hoechst solution 1:3000 were applied at room temperature for 1 h. Washing was carried out 3 times for 5 minutes at room temperature with PBS and stained coverslips were mounted in fluoromont solution (ThermoScientific) and imaged using a LSM-800 confocal microscope (Zeiss) with Zen software. Images were exported, analyzed and processed with Fiji software.

### RFP-GFP-LC3B tandem sensor assay

Patient’s fibroblasts were seeded on coverslips coated with 50μg/ml of collagen I. After 24h, cells were transfected with Premo™ autophagy tandem sensor RFP-GFP-LC3B kit (ThermoScientific) according to the manufacturer’s protocol. After 24h, cells were treated with 10 μM givinostat or 0.1% DMSO. After 24h of treatment, cells were fixed with 4% formaldehyde and counterstained with Hoechst 33342 solution (Invitrogen). Stained coverslips were mounted in fluoromont solution and imaged using a LSM-800 confocal microscope.

### Transmission electron microscopy

R77C-α-SGmCh fibroblasts seeded on coverslips coated with 50 μg/ml of collagen were chemically fixed in 2.5 % (v/v) glutaraldehyde, 2 % (v/v) paraformaldehyde in 0.1 M cacodylate (pH 7.2) buffer for 2h at room temperature, washed in cacodylate (0.1 M), post-fixed with 2 % (w/v) osmium tetroxide supplemented with 1.5 % (w/v) potassium ferrocyanide (45 min, 4°C), washed in water, dehydrated in ethanol (increasing concentration from 30 to 100 %) and embedded in Epon as described in (Hurbain et al., 2017). Ultrathin sections of cell monolayers were prepared with a Reichert Ultracut S ultramicrotome (Leica Microsystems) and contrasted with uranyl acetate and lead citrate. Electron micrographs were acquired on a Tecnai Spirit Electron Microscope (ThermoFischer Scientific) equipped with a 4k CCD camera (Quemesa, EMSIS GmbH, Muenster, Germany) using iTEM software (EMSIS).

### YFP-based CFTR functional assay

The assay was performed according to (Galietta et al., 2001). Briefly, CFTR activity was measured in stably transfected HEK cells coexpressing the halide-sensitive yellow fluorescent protein YFP-H148Q/I152L and F508del CFTR mutant. Cells were seeded in poly-l-lysine coated 96-well black/clear bottom microplates. The CFTR functional assay was carried out 24 hours after individual compound treatments with 3 μM VX-809, 10 μM givinostat, 5 nM BTZ or combined treatments. Cells were incubated for 30 min with 50 μl of PBS containing CPT-AMPc (100 μM) before to be transferred to a TrisStar plate reader (Berthold). When indicated, before being treated with of PBS containing CPT-AMPc, cells were incubated with 10 μM VX-770 for 30 min. Each condition was tested in triplicate. Cell fluorescence (excitation: 485 nm; emission: 535 nm) was continuously measured before and after addition of 100 μl of PBS-NaI. Cell fluorescence recordings were normalized to the initial average value measured and signal decay was fitted to an exponential function to derive the maximal slope. Maximal slopes were converted to rates of change in intracellular I-concentration (in mM/sec). The experiments were repeated three times.

### Statistical analysis

Data are presented as means ± SD. Statistical analysis was performed using the Student’s t test. Statistical significance was considered for ***P≤0.001; **P≤0.01; *P≤0.05. Curvefitting, histograms, IC_50_ and EC_50_ determinations were performed using GraphPad Prism (v8.4.3).

## Results

### Identification of HSP90 and HDAC families as new inhibitors of misfolded R77C-α-SG protein degradation using combinatorial high-content screening

This study was performed on immortalized fibroblasts from a LGMD R3 patient homozygous for R77C-α-SG mutation and stably transduced with a R77C-α-SGmCh reporter lentivirus. This model was previously described as a relevant cellular model to concomitantly investigate the R77C-α-SG expression and localization after compound treatments (Hoch et al., 2019). The drug screening strategy is based on simultaneous identification of positive cells for heterologous R77C-α-SGmCh protein by immunostaining and for presence of the R77C-α-SGmCh protein at the cell membrane compartment using a high-content imaging automate in 384 well plates on non-permeabilized cells **(Supplementary Figure S1A**). Masks and algorithms were used to define the nuclear (Hoechst), membrane α-SG staining and mCherry fluorescent signal **(Supplementary Figure S1B**). First, dose response experiments were carried out with BTZ to identify the toxicological profile (**Figure 1A**) as well as IC_50_ on proteasome activity and EC_50_ on R77C-α-SG membrane rescue (**Figure 1B**). Analysis of these data allowed us to identify a non-toxic dose of BTZ (5 nM) having limited effect on R77C-α-SG membrane rescue and partial inhibition of the proteasome activity. In comparison, 30 nM BTZ was identified as a dose highly rescuing the R77C-α-SG membrane expression with a complete inhibition of the proteasome activity (**Figures 1B,C**). Accordingly, a combinatorial screening of a chemical library of 958 FDA-approved drugs at 5 μM was performed in the presence of 5 nM BTZ (**Figure 1D**), hypothesizing that the partial proteasome inhibition would permit to identify drugs acting in synergy. A simultaneous screening of the chemical library alone at 5 μM was performed as a control (**Figure 1E**). Quantification of double positive cells for the negative and positive controls (0.1% DMSO and 30 nM BTZ, respectively) of each plate of the screening was carried out **(Supplementary Figure S1C**), confirming a statistical relevant difference between controls with a mean Z’ factor of 0.5 **(Supplementary Figure S1D**). Compounds were considered as potential candidates when their effect of directing the mutated protein at the membrane was superior to two standard deviations from the mean of all the tested compounds, and their cell viability was superior to 45%. This led to an initial list of 31 hits when the library was tested alone and 39 hits when tested in combination with 5 nM BTZ with an overlap of 23 hits between the two conditions and 16 unique to the combination (**Figure 1F and Supplementary Table S2**). Interestingly, among the list of 39 hits, 9 HSP90 and 8 histone deacetylase HDAC inhibitors were identified as significantly efficient on R77C-α-SG membrane expression (**Figures 1D,E and Supplementary Table S2**). The identification of these two families of compounds highlights new potential pathways implicated in the degradation of the R77C-α-SG protein. Interestingly, previous studies described a link between HDACs and HSP90 signaling pathways as HDAC6 regulates HSP90 deacetylation, an event critical for maintaining its normal chaperone activity (Bali et al., 2005; Kovacs et al., 2005).

**Figure 1.**
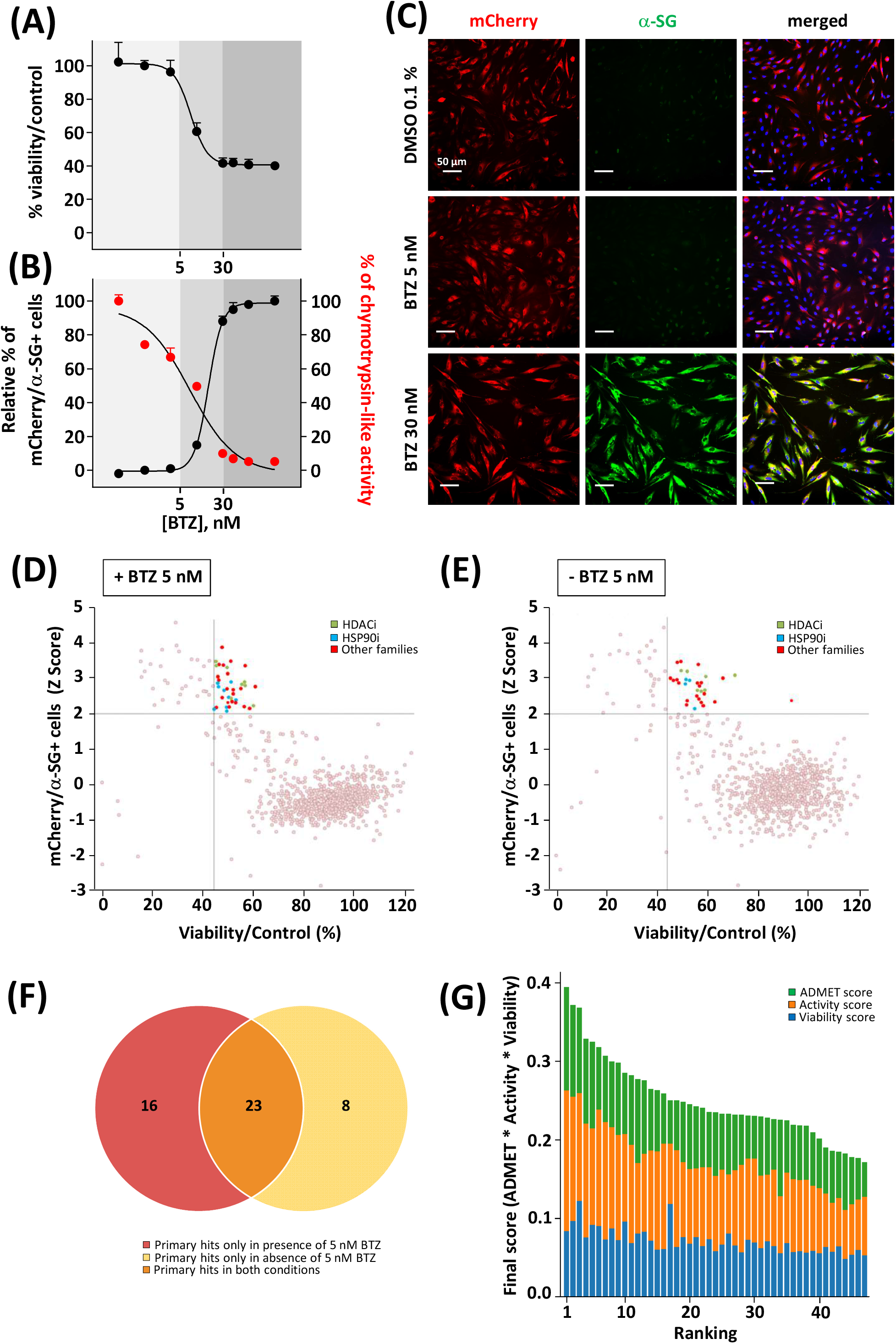
Combinatorial high-content screening for R77C-α-SGmCh membrane rescue. **(A-B)** Cell viability **(A)**, chymotrypsin-like activity of the proteasome and quantification of mCherry and membrane α-SG positive fibroblasts **(B)** following treatment with increasing concentrations of BTZ. Each point represents the mean±SD (n=4) of a representative experiment over 3 independent experiments. **(C)** mCherry fluorescent signal (red) and α-SG staining (green) in non-permeabilized condition in fibroblasts overexpressing R77C-α-SGmCh treated with 0.1% DMSO, 5 nM or 30 nM BTZ. Nuclei are labelled by Hoechst staining (blue). Scale bar = 50 μm. **(D-E)** Dot plot representations of the effects of the 958 drugs at 5 μM in combination with 5 nM **(D)** or alone **(E)** on R77C α-SGmCh membrane expression (Z score >2) and cell viability (viability >45%). Hits are colored according to their compound families, HDACi are green, HSP90i are blue and other families are red. **(F)** Venn diagram illustrating the number of primary hits only in presence of 5 nM BTZ (red), only in absence of 5 nM BTZ (yellow) or in both conditions (orange). **(G)** Classification of hit compounds based on a virtual ADMET analysis using artificial intelligence. The final score comprised the ADMET score (green), the activity score (orange) and the viability score (blue) determined during the screening. BTZ = bortezomib.

### Investigation of givinostat for R77C-α-SG membrane rescue

We then performed a virtual Absorption, Distribution, Metabolism, Elimination and Toxicity (ADMET) analysis of the hit compounds using artificial intelligence, evaluating 76 properties. The compound ranking metric used is an equally balanced weight of the percentage of ADMET endpoints passing, among which 49 machine learning predictions, the cell viability and the compound activity on the bioassay. While in this classification, givinostat was ranked in the top 10 compounds (**Figure 1G and Supplementary Table S3**), the review of various factors of the top compounds such as their clinical approval stage, their indications, and their targets, highlighted givinostat as one of the most promising orphan drugs to be investigated for its effect on α-SG. Besides its good safety profile, givinostat, retained our attention because of its anti-fibrotic effect in Duchenne muscular dystrophy (DMD) animal models (Consalvi et al., 2013) and its current evaluation in a phase III clinical trial in DMD patients. Doseresponse experiments for membrane rescue of R77C-α-SGmCh were conducted with increasing concentrations from nanomolar to micromolar range of givinostat alone or in combination with 5 nM BTZ. Analysis of α-SG staining following givinostat treatment alone revealed a punctiform α-SG signal suggesting a partial rescue of the protein degradation (**Figure 2A, left panels**). However, similar experiment conducted in presence of the combined treatment of givinostat with 5 nM BTZ revealed a continuous α-SG signal at the membrane compartment (**Figure 2A, right panels**). These observations were confirmed by performing a dose-reponse experiment quantifying the presence of R77C-α-SGmCh at the membrane that revealed a significantly higher efficacy of the combined treatments (EC_50_ of 0.8 μM) as compared to givinostat treatment alone (EC_50_ of 2.5 μM) (**Figure 2B**) without additional toxicity (**Figure 2C**). α-SG protein level was then investigated by immunoblot analysis in presence of the different treatments (**Figure 2D**), revealing an increased α-SG protein level after treatment with 10 μM givinostat in both presence and absence of 5 nM BTZ. This result suggested that givinostat treatment alone could induce the inhibition of the R77C-α-SG protein degradation but probably not its full trafficking to the membrane as indicated by the imperfect immunostaining at the membrane (**Figure 2A**). Of note, as previously reported by our group, treatment with 30 nM BTZ leads to the appearance of an extra band corresponding to an immature form of α-SG (Hoch et al., 2019). This additional band was not observed in presence of 10 μM givinostat alone, suggesting a distinct mechanism of action of both compounds. Interestingly, similar results were obtained with parallel experiments following belinostat treatment, another pan-HDACi identified in the screening, alone or in combination with 5 nM BTZ, suggesting a common mode of action of these two HDACi for the R77C-α-SG membrane expression rescue **(Supplementary Figure S2**). To explore givinostat mode of action on α-SG, we next evaluated the endogenous and exogenous *SGCA* gene expression after treatments with 10 μM givinostat or 30 nM BTZ (**Figure 2E**). Analysis of qPCR revealed that gene expression was not increased by any treatment suggesting that R77C-α-SG protein rescue was not due to *SGCA* transcriptional increase. Inversely, we observed a significant transcriptional decrease that did not reflect the increase of R77C-α-SG protein level, suggesting a post-transcriptional effect of givinostat on R77C-α-SG protein.

**Figure 2.**
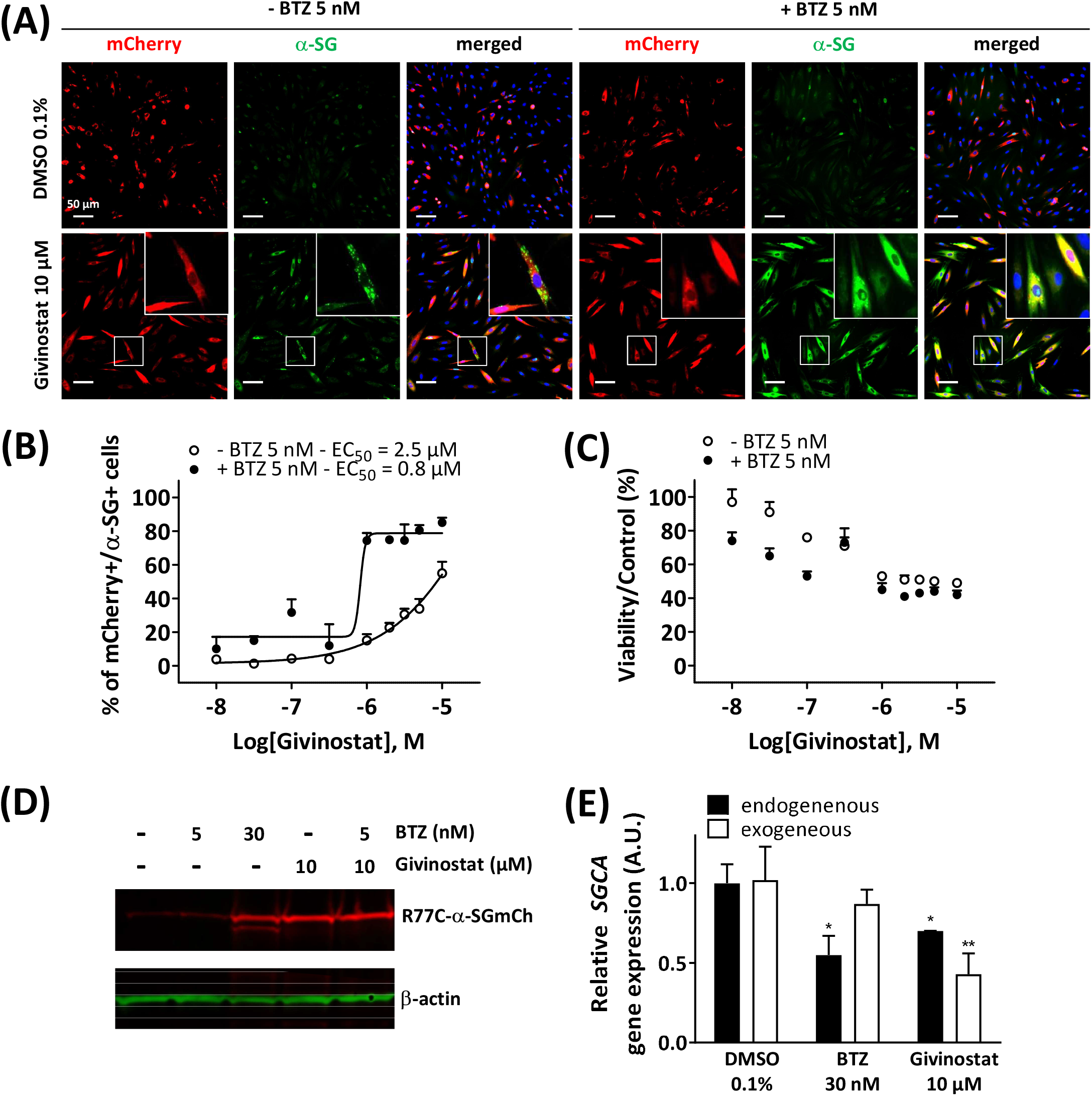
Rescue of misfolded R77C-α-SG degradation by givinostat. **(A)** Images of mCherry fluorescent signal (red) and membrane α-SG staining under non-permeabilized condition (green) in fibroblasts overexpressing R77C-α-SGmCh and treated with DMSO (0.1%), BTZ (5 nM or 30 nM) or givinostat (10 μM or increasing concentrations) in absence or in presence of 5 nM BTZ for 24 hours. Nuclei are labelled by Hoechst staining (blue). Scale bar = 50 μm. **(B-C)** Quantification of mCherry and membrane α-SG positive fibroblasts **(B)** and cell viability **(C)** in fibroblasts. Each point represents the mean±SD (n=4) of a representative experiment over 3 independent experiments. **(D)** Immunoblot analysis of α-SG expression after fibroblasts treatments. ß-actin was a loading control. **(E)** Measure of endogenous and exogenous *SGCA* gene expression by qPCR following fibroblasts treatments. Values are expressed as relative *SGCA* gene expression to DMSO treated cells. Data are mean±SD (n=3) of a representative experiment over 3 independent experiments, *P≡0.05 (Student’s t-test). BTZ = bortezomib.

### Rescue of different missense mutations of SG proteins through combination of givinostat and bortezomib treatments

To further validate the interest of givinostat, we evaluated the effect of this compound alone or combined with BTZ on other missense mutations leading to misfolded α-SG and γ-SG and frequently reported in LGMD R3 and LGMD R5 patients. Lentiviruses expressing the V247M, R34H or I124T α-SGmCh or the C283Y or E263K γ-SGmCh mutants were transduced in immortalized R77C patient fibroblasts or SGCG^-/-^ fibroblasts, respectively. Immunofluorescence images analysis in non-permeabilized condition revealed the absence of detectable α-SG or γ-SG staining at the membrane level in all mutant transduced cells (**Figure 3A and Supplementary Figure S3A**). Analysis of α-SG and γ-SG mutant proteins localization was then evaluated following treatment with BTZ (5 nM and 30 nM) or givinostat (10 μM alone or combined with 5 nM BTZ). All mutant proteins were rescued with 30 nM BTZ, confirming that these misfolded proteins are rescuable from proteasomal degradation. Combination of givinostat and BTZ treatments leads to the rescue of all mutated protein, except the I124T-α-SGmCh, with different efficacies ranging between 40 % and 60 % of rescued cells (**Figure 3B and Supplementary Figures S3C,D**).

**Figure 3.**
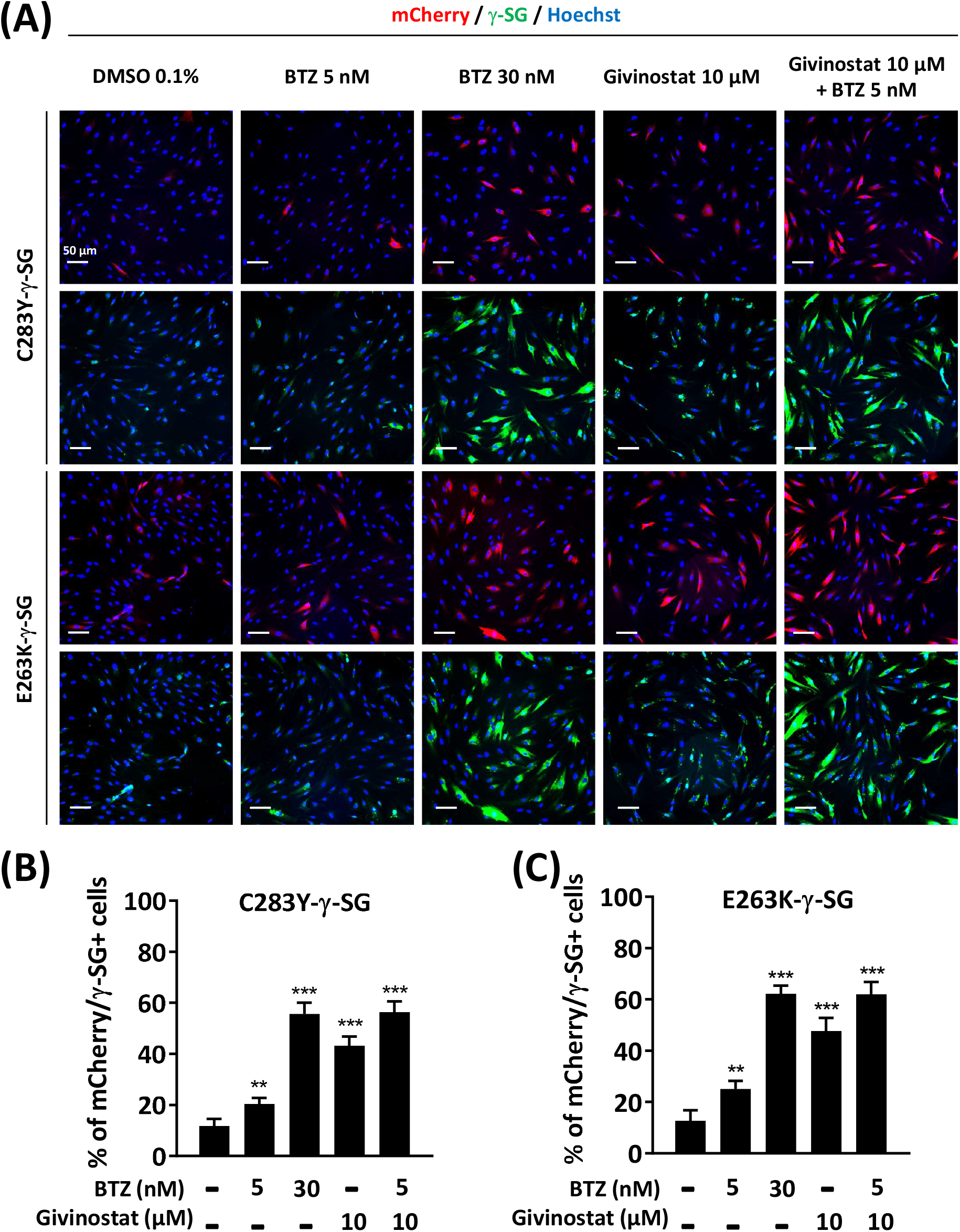
Evaluation of givinostat and bortezomib combination on γ-SG mutated proteins. **(A)** Images of mCherry fluorescent signal (red) and membrane γ-SG staining under non-permeabilized condition (green) of SGCG−/− fibroblasts transduced with lentivirus expressing C283Y-γ-SGmCh or E263K-γ-SGmCh constructs and treated with DMSO (0.1%), BTZ (5 nM and 30 nM) or givinostat (10 μM) in presence or in absence of 5 nM BTZ for 24 hours. Nuclei are labelled by Hoechst staining (blue). Scale bar = 50 μm. **(B-C)** Quantification of mCherry and membrane C283Y-γ-SG **(B)** and E263K-γ-SG **(C)** positive fibroblasts. Each chart represents the mean±SD (n=4) of a representative experiment over 3 independent experiments; *P≡0.05, ***P≤0.001 (Student’s t-test). BTZ = bortezomib.

### Givinostat treatment corrects F508del CFTR trafficking

Recent evidences have reported the role of HDAC inhibition in the transport at the plasma membrane of the misfolded F508del-CFTR protein, the most frequent mutant causing cystic fibrosis (CF) (Anglès et al., 2019). CF is an inherited disorder that causes severe damage to the lungs, digestive system and other organs in the body. It is now well established that restoring the expression of the F508del-CFTR mutant on the surface of epithelial cells by inhibiting its degradation allows the correction of CF patients phenotype since the F508del-CFTR protein is partially functional when it is correctly positioned at the apical pole of the cells. To open up the possibility of using givinostat for other misfolded protein rescue, we investigated the impact of givinostat alone or in combination with the FDA-approved CF treatments Lumacaftor (VX-809) or Orkambi (VX-770 + VX-809) on CFTR (Van Goor et al., 2006, 2009). To do so, we used embryonic kidney (HEK293 MSR Grip Tite) cells stably coexpressing a yellow fluorescent protein (eYFP) variant and F508del-CFTR-3HA (Trzcińska-Daneluti et al., 2015). Because the fluorescence of eYFP variant can be quenched in response to iodide influx entering the cell through a functional CFTR channel, this strategy is used to indirectly measure CFTR activity. Results of this experiment revealed that treatment with 10 μM givinostat induced a similar level of eYFP quenching to the one observed with 3 μM Lumacaftor (**Figures 4A and 4C**). Importantly, an additive effect on the CFTR channel activity was observed using the combination of givinostat with Lumacaftor treatment (**Figures 4A and 4C**) and synergistic effects were observed with Orkambi treatment (**Figures 4B and 4D**). Results were confirmed by measuring CFTR protein levels (B- and C-band) following treatments with 10 μM givinostat, 3 μM Lumacaftor or combinational treatments by immunoblot analysis showing that givinostat induces a synergistic improvement in the trafficking efficiency of the F508del CFTR mutant **(Figures 4E-G**).

**Figure 4.**
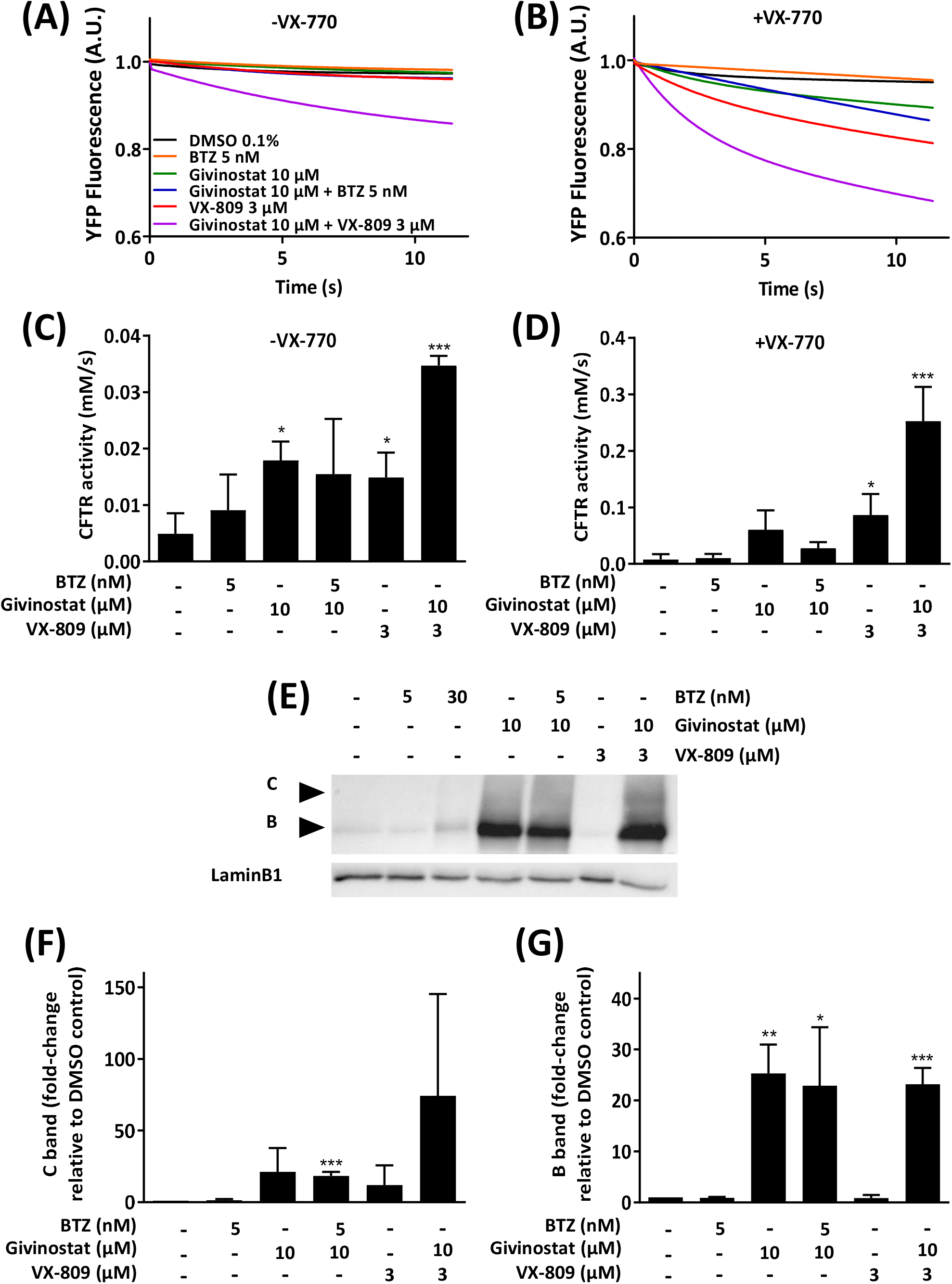
Evaluation of givinostat on CFTR activity. **(A-D)** Measure of YFP fluorescence **(A, B)** and quantification of CFTR activity **(C, D)** in HEK-F508del/YFP cells treated with different combinations of BTZ (5 nM), givinostat (10 μM) or VX-809 (3 μM) for 24 hours in absence **(A, C)** or in presence **(B, D)** of VX-770 (10 μM) for 30 min. **(E-G)** Immunoblot analysis **(E)** and quantification of the C-band **(F)** or the B-band **(G)** of CFTR expression in CFBE-F508del cells treated with different combinations of BTZ (5 nM), givinostat (10 μM) or VX-809 (3 μM) for 24 hours. LaminB1 was used to evaluate the loading. Values are expressed as fold-change relative to DMSO 0.1% treatment. Data are the mean±SD of 3 independent experiments; *P≤0.05, **P≤0.01, ***P≤0.001 (Student’s t-test). BTZ = bortezomib.

### Givinostat treatment induces acetylation of HSP90 and α-tubulin

Following these demonstrations of the interest of givinostat in various pathological conditions, we explored its mechanism of action. First and because we identified several HSP90i in the combinatorial screening hits list suggesting the involvement of HSP90 chaperone activity in the degradation of the R77C-α-SG protein and considering previous studies already connecting HDACs and HSP90 signaling pathways (Rao et al., 2012), we investigated the effect of givinostat on HSP90. HDACs function through the removal of acetyl groups attached to lysine residues in histones and non-histone proteins. Interestingly, it has been previously demonstrated that at least one HDAC, HDAC6, functions as an HSP90 deacetylase (Kovacs et al., 2005) with the inactivation of HDAC6 leading to HSP90 hyperacetylation and resulting in a reduced interaction with client proteins (Bali et al., 2005). After treatment of fibroblasts with 10 μM givinostat, immunoprecipitation of HSP90 followed by immunoblotting with antibodies recognizing acetylated lysine residues and α-SG proteins were performed. An important acetylation of HSP90 but no effect on the levels of the protein was observed after treatment (**Figure 5A**). Interestingly, α-SG was identified to interact with HSP90 but the givinostat treatment did not alter this interaction (**Figure 5B**), suggesting that the rescuing effect of givinostat on α-SG does not implicate a reduction of α-SG as a client protein of HPS90.

**Figure 5.**
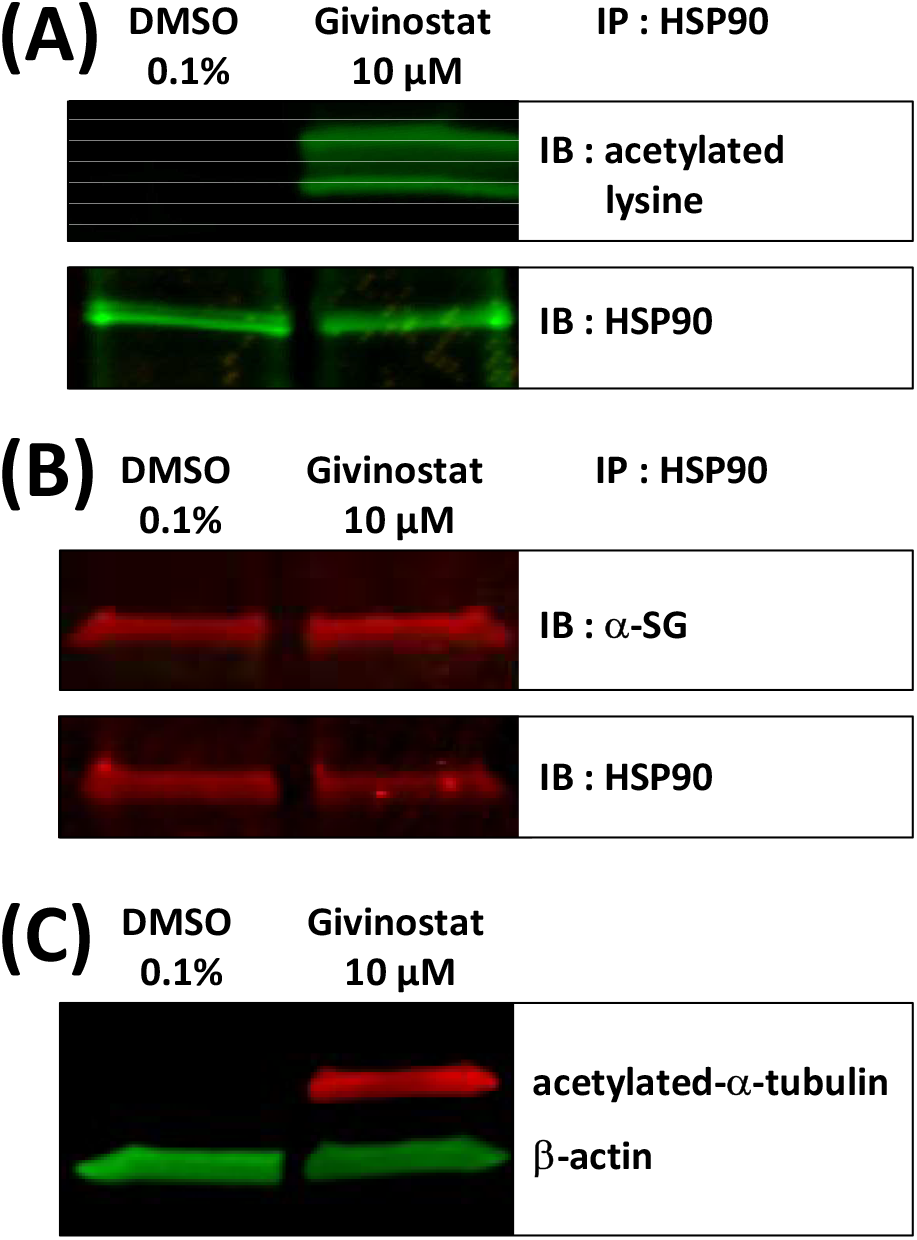
Givinostat regulates HSP90 and α-tubulin acetylation. **(A-B)** Total cell lysates of fibroblasts overexpressing R77C-α-SGmCh treated with givinostat (10 μM) or DMSO (0,1%) for 24 hours were prepared and immunoprecipitated to HSP90. Immunoblot analysis was performed with anti-acetylated-lysine **(A)**, with anti-α-SG **(B)** and anti-HSP90 antibodies. **(C)** Immunoblot analysis acetylated-α-tubulin expression in treated fibroblasts. β-actin was a loading control.

After confirmation that givinostat modulates the acetylation of HSP90, the hypothesis of the implication of HDAC6 was then investigated. To validate the modulation of HDAC6 activity following givinostat treatment, we analyzed by immunoblot the acetylation of α-tubulin, an event mostly dependent on HDAC6 activity. As hypothesized, givinostat treatment induced the acetylation of α-tubulin (**Figure 5C**). We then examined the effect of tubacin, a selective HDAC6 inhibitor, on R77C-α-SGmCh mutant rescue. Our results demonstrated that tubacin induced a dose-reponse curve for membrane R77C-α-SGmCh expression and a synergistic effect with 5 nM BTZ **(Supplementary Figures S4A,B**) with cell toxicity in the same range than givinostat **(Supplementary Figure S4C**). Immunoblot analysis confirmed the increase of α-SG protein levels and the acetylation of α-tubulin **(Supplementary Figure S4D**) following combined tubacin and BTZ treatment. These data suggest that the inhibitory effect of givinostat leading to the R77C-α-SG protein rescue could implicate the HDAC6 activity.

### Givinostat mode of action is not occurring at the proteasome level

As a second step, we analysed givinostat effect on degradative systems, starting by the proteasome activity considering the synergistic effect of the BTZ and givinostat combination. To do so, we first measured the chymotrypsin-like activity of the proteasome after treatment with increasing concentrations of givinostat (**Supplementary Figure S5A**). Results revealed no significant decrease of the chymotrypsin-like activity following givinostat treatment, suggesting a distinct mechanism of action than BTZ. The same conclusion was obtained with parallel experiments following belinostat treatment **(Supplementary Figure S2E**). We then evaluated the effect of givinostat on the BTZ-mediated proteasome inhibition by measuring the chymotrypsin-like activity after treatments with increasing concentrations of BTZ in the presence of 10 nM, 30 nM, 100 nM, 300 nM, 1 μM, 3 μM or 10 μM of givinostat (**Supplementary Figure S5B**). Analysis of the dose-response curves revealed no significant IC_50_ changes of the chymotrypsin-like activity induced by the different combinations of BTZ and givinostat, while significant EC50 changes of the membrane R77C-α-SGmCh mutant protein rescue were observed (**Supplementary Figures S5C-E**), providing evidences that givinostat had no effect on BTZ-mediated proteasome inhibition, although toxicity is observed at high dose combinations. Similar results were obtained by testing the combination of givinostat with two other proteasome inhibitors respectively, Carfilzomib (CFZ) and MG132 **(Supplementary Figure S6**). As previously done with BTZ, quantification of the percentage of cells with membrane localization of R77C-α-SGmCh after treatment with increasing concentrations of givinostat combined to low doses of CFZ (10 nM) or MG132 (100 nM) indicated their synergistic effect on R77C-α-SGmCh mutant protein rescue **(Supplementary Figure S6G**).

### Givinostat treatment inhibits autophagy at the late stage

To deeper investigate the mechanism of givinostat on membrane SG rescue, we analyzed the ubiquitination of proteins. Immunoblot analysis revealed similar increased expression levels of ubiquitinated proteins after 10 μM givinostat or 5 nM BTZ treatment and additive effects were observed with combination of both treatments (**Figure 6A, upper panel**). These results, together with the fact that givinostat did not inhibit the proteasome activity nor act through increasing the effect of BTZ on proteasome inhibition, suggested that another mechanism of protein degradation should be prevented by the effect of givinostat. Beside the ERAD pathway, another important degradation pathway for proteins present in the ER is ER-phagy, a selective form of autophagy which enables the formation of autophagosomes from the ER membranes and mediates their fusion with lysosomes for proteins degradation (Song et al., 2018). Interestingly, it was previously described that R77C-α-SG proteins can form aggresomes in the ER, an element compatible with the possibility of activation of ER-phagy with mutated sarcoglycans (Draviam et al., 2006). In line with these results, mCherry aggregates were observed by confocal immunofluorescence analysis in the ER after 30 nM BTZ treatment **(Supplementary Figure S7**) revealing R77C-α-SG proteins aggregation when the proteasome activity is inhibited. In comparison, no aggregates were observed after 10 μM givinostat treatment **(Supplementary Figure S7**). Since recent studies have demonstrated that proteasome inhibitors could induce autophagy as a compensatory response (Kocaturk and Gozuacik, 2018), we hypothesized that BTZ treatment, in addition to its inhibitory effect on proteasome, would orient R77C-α-SG proteins towards the autophagic route. Our observation of the synergistic effects of givinostat when associated to BTZ led to the hypothesis that givinostat effect on R77C-α-SG proteins was relied on the regulation of this alternative autophagic route. We investigated the autophagy process by monitoring the conversion of LC3B from the cytosolic form, LC3B-I, to the membrane-associated form, LC3B-II (Kabeya et al., 2000) using immunoblot analysis. Conversion to LC3B-II was induced in dose-dependent manner in cells upon BTZ treatment confirming that autophagy was activated in a compensatory response to proteasome inhibition (**Figures 6A, lower panel, and 6B**). Then, we assessed the expression of P62, another marker of autophagy selectively incorporated into autophagosomes through direct binding with LC3B. It has also been reported that P62 is degraded at the late stage of autophagy process, and that accumulation of P62 could be the consequence of an autophagic flux inhibition meaning that autophagy is activated but not completed (Mizushima et al., 2010). Upon 10 μM givinostat treatment, accumulation of P62 and LC3B-II were observed suggesting an inhibition of the autophagic flux (**Figures 6A, lower panel, and 6B,C**). Interestingly, P62 accumulation was also observed upon BTZ treatment (**Figures 6A, lower panel, and 6C**). In line with these results, many studies have reported that P62 transcription was rapidly upregulated upon proteasome inhibition (Sha et al., 2018). To further understand the mechanism of autophagy inhibition induced by givinostat, we next explored the process of autophagy by immunofluorescence imaging using a RFP-GFP-LC3B tandem sensor allowing the dissection of the maturation of autophagosomes to functional autolysosomes. By combining an acid-sensitive GFP-LC3B sensor with an acid-insensitive RFP-LC3B sensor, the transition from neutral autophagosomes to acidic autolysosomes can be visualized by imaging the specific loss of the GFP fluorescence in cells. Patient’s fibroblasts were transiently transduced with a RFP-GFP-LC3B tandem sensor and treated with 10 μM givinostat or 0.1% DMSO. Givinostat treatement dramatically increased the number of RFP-LC3B structures positive for GFP signal (**Figure 6D**) indicating an accumulation of non-acid autophagic structures. Also, LAMP2 staining was largely increased after givinostat treatment (**Figure 6E**) suggesting that the generated autophagic structures were positive for lysosomal components. To decipher whether givinostat affects autophagosome-lysosome fusion or lysosomal pH increase, we examined the ultrastructure of the degradative organelles by using transmission electron microscopy (TEM). Accordingly, and as compared to controls, givinostat-treated cells showed a dramatic accumulation of large and irregular electron-dense single membrane-bound organelles containing a large amount of luminal proteinaceous and lipid materials (**Figures 7A, white arrows, and 7B)**. Those organelles were not autophagosomes, which were equally identified in DMSO- and givinostat-treated cells as electron-lucent organelles delimited by a double membrane (**Figures7A, black arrows, and 7B)** suggesting similar autophagy induction with both treatments. While those organelles shared ultrastructural features of autolysosomes and would evidence that autophagosomes fused with lysosomes, the nondegraded luminal content was suggestive of a lower degradative capacity of these structures upon givinostat treatment.

**Figure 6.**
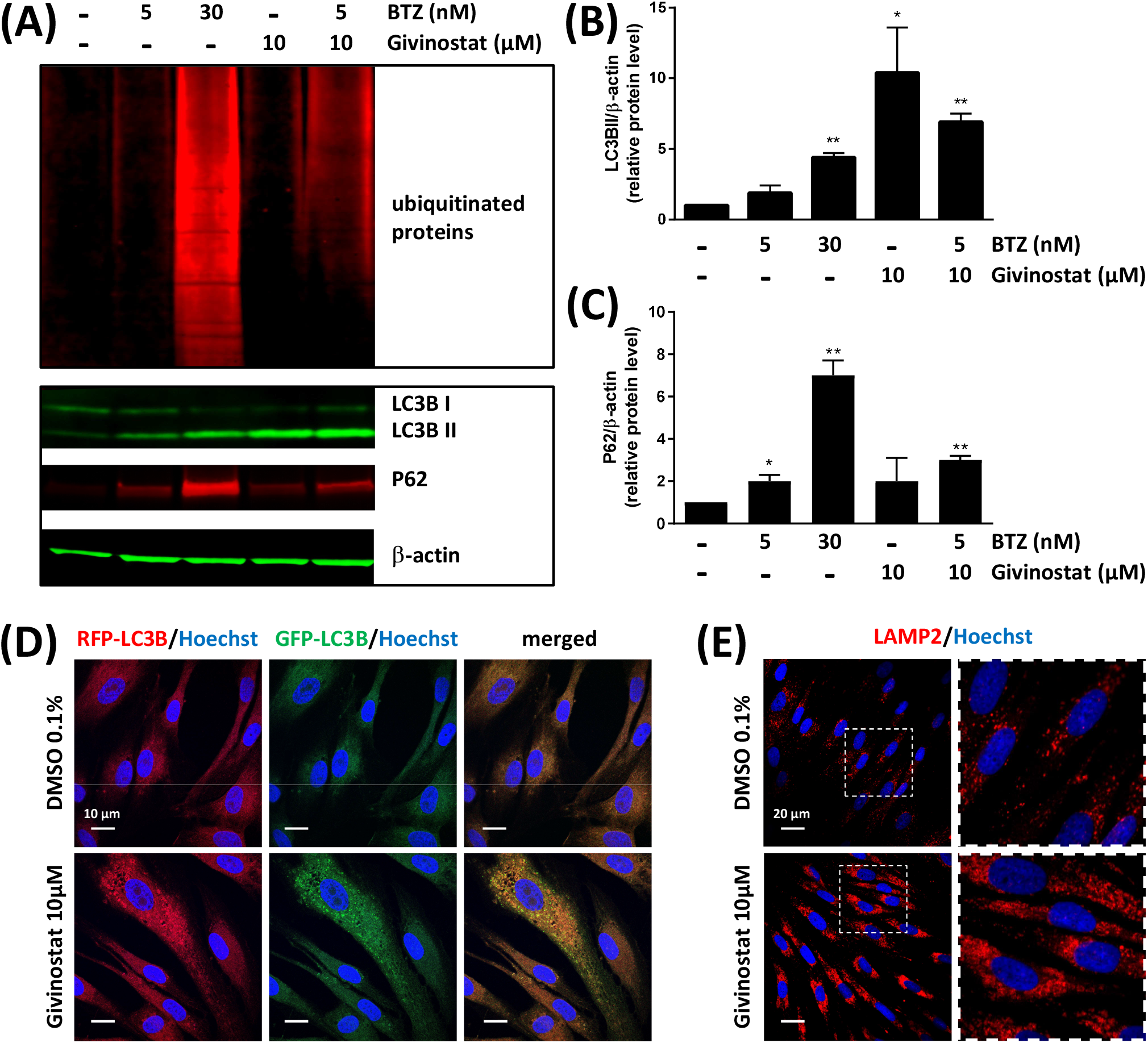
Givinostat inhibits the autophagy pathway. **(A)** Immunoblot analysis of ubiquitinated proteins (upper pannel), LC3BI / II and P62 (lower panel) expression in fibroblasts overexpressing R77C-α-SGmCh and treated with DMSO (0.1%), BTZ (5 nM and 30 nM) or givinostat (10 μM) in absence or in presence of 5 nM BTZ for 24 hours. β-actin was used to evaluate the loading. **(B-C)** Quantification of LC3BII **(B)** and P62 **(C)** protein expression levels. Values are expressed as relative protein expression level to DMSO treated cells and normalized to β-actin. Data are mean±SD of 3 independent experiments; *P≤0.05, **P≡0.01 (Student’s t-test). **(D)** Fibroblasts transiently expressing GFP-RFP-LC3B were treated with givinostat (10 μM) or DMSO (0,1%) for 24 hours and the colocalization of GFP and RFP puncta was detected. Nuclei are labelled by Hoechst staining (blue). Scale bar = 10 μm **(E)** Confocal images of LAMP2 (red) and Hoechst staining (blue) after DMSO (0.1%) or givinostat (10 μM) treatments. Scale bar = 20 μm. Insets are magnifications of the boxed areas.

**Figure 7.**
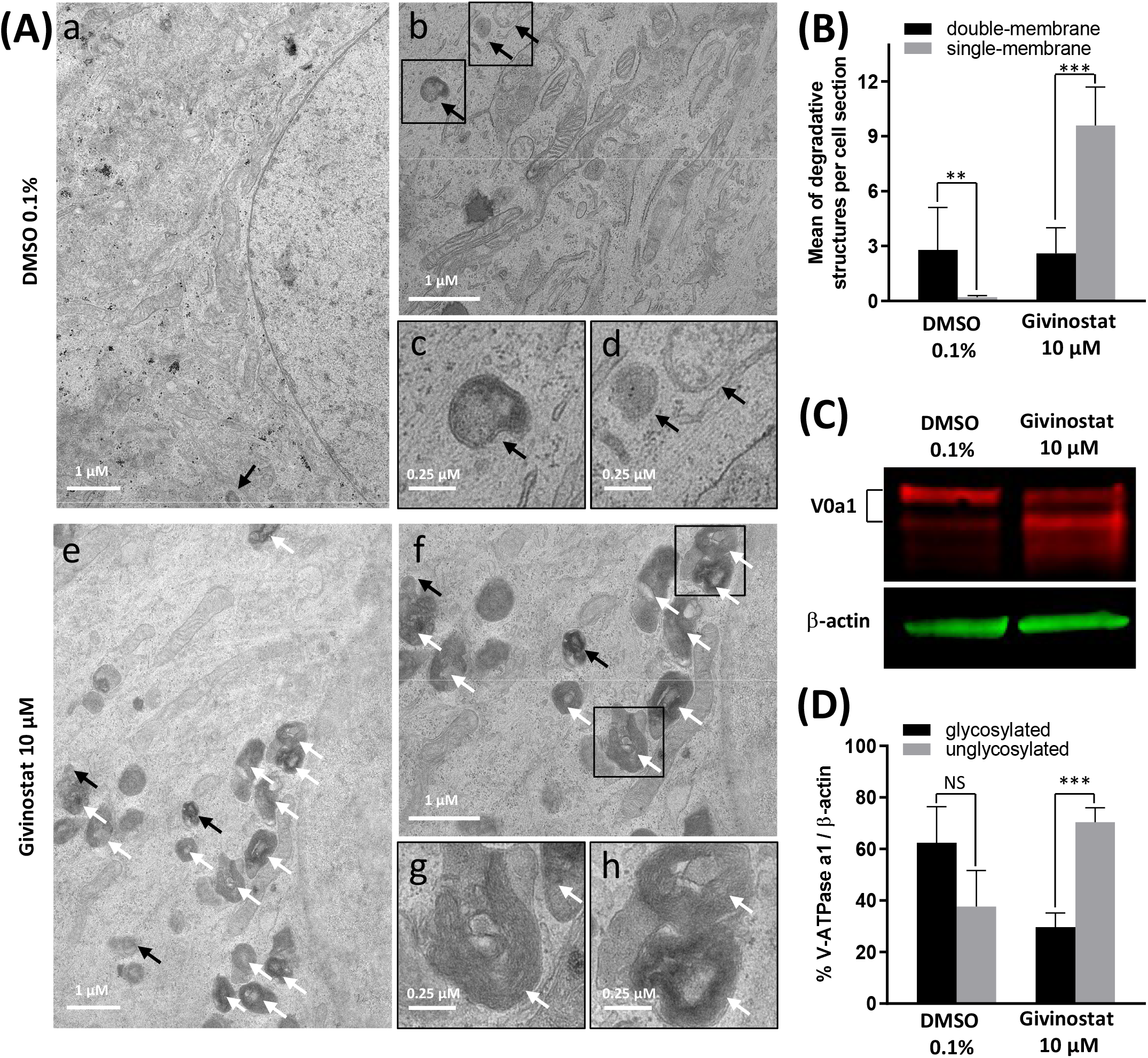
Givinostat impairs autolysosomes degradation. **(A)** Transmission electron microscopy analysis of fibroblasts overexpressing R77C-α-SGmCh and treated with DMSO (0.1%) or givinostat (10 μM) for 24 hours. Few double membrane-bound organelles with a relative electron-lucent luminal content (black arrows) were found in DMSO and givinostat treated cells. Givinostat induced a large accumulation of single membrane-bound organelles with electron dense lumen (white arrows). Insets are magnifications of the boxed areas. **(B)** Quantification of degradative structures identified according to their morphology described in A. Data are mean±SD of degradative structures identified per cell section (n=7); *P≤0.05, ***P≤0.001 (Student’s t-test). **(C)** Immunoblot analysis of the v-ATPase V0a1 subunit expression in fibroblasts treated with DMSO (0.1%) or givinostat (10 μM) for 24 hours. β-actin was used to evaluate the loading. **(D)** Quantification of the glycosylated and the unglycosylated V0a1 subunit expression. Values are expressed as percentage of total V0a1 protein expression normalized to β-actin. Data are mean±SD of 3 independent experiments; NS=non-significant, ***P≤0.001 (Student’s t-test).

### Givinostat treatment inhibits the N-glycosylation of the V-ATPase V0 subunit a1

The degradative function of lysosomes is carried out by hydrolytic enzymes within their lumen that require acidic pH for optimal activity (Mellman et al., 1986). Acidification of lysosomes is mainly controlled by the vacuolar-type ATPase (V-ATPase), a multimeric protein composed of two subcomplexes, the cytosolic V1, and the membrane-bound V0, allowing transport of protons across the lysosomal membrane. Reversible assembly of V1/V0 sub-complexes on lysosomal membrane regulates the proton pump activity of the V-ATPase. To probe for abnormal lysosomal acidification, we investigated the N-glycosylation of the V-ATPase subunit V0a1, which has been demonstrated to regulate the proton pump assembly in lysosomes (Lee et al., 2010a). Immunoblot analysis of fibroblasts treated with 10 μM givinostat or 0.1% DMSO using an anti-V-ATPase V0a1 antibody revealed two bands corresponding to the non-glycosylated (100 kDa) and the mature N-glycosylated (120 kDa) subunit V0a1. Fibroblasts treated with 10 μM givinostat highly expressed nonglycosylated V0a1 in comparison to 0.1% DMSO treated cells which expressed mainly mature N-glycosylated V0a1 (**Figures 7C,D**). These data strongly support the conclusion that, under 10 μM givinostat treatment, the V-ATPase V0a1 subunit is not N-glycosylated, preventing acidification of lysosomes and therefore degradation of autolysosome content.

## Discussion

The main result of this study is the identification of the synergistic effect of givinostat with a low dose of BTZ on misfolded protein degradation through the dual blockade of autophagy and proteasome pathways. Overall our results describe a new therapeutic avenue for the treatment of LGMD R3 patients carrying missense mutations of *SGCA* but also other genetic diseases sharing similar protein degradation defects as LGMD R5 and CF.

Proteasome inhibitors are mainly used for the treatment of multiple myeloma and lymphoma mostly because of their effects on cell cycle and apoptosis (Fricker, 2019). To date, BTZ is the most used FDA-approved proteasome inhibitor. While efficient on tumors growth, BTZ treatment is used for short term therapy because of its well reported side effects (Kaplan et al., 2017; Mohan et al., 2017) that preclude its use for long term administration in patients. Here, we report that α-SG membrane rescue induced by BTZ arise only at toxic doses inhibiting proteasome up to 95% whereas conversely, proteasome inhibition inferior or equal to 50% does not lead to any effect on α-SG. Based on these observations, we used HTS to evaluate the effect of 958 FDA-approved drugs and identify those capable to increase R77C-α-SG rescue without increasing BTZ-induced proteasome inhibition and its associated cell toxicity. In the past decade, similar strategies have already been reported to identify drugs increasing effects (Chumakov et al., 2014; Yu et al., 2019) or alleviating side effects (Hajj et al., 2015; Zhou et al., 2019) of other therapeutic agents. In this study, we identified 47 compounds all capable to increase R77C-α-SG rescue and belonging mostly to HSP90i, HDACi, aurora kinase inhibitors (AKi) and cyclin-dependent kinase inhibitors (CDKi) families. Among this hits list, although having an effect on R77C-α-SG rescue, 16 compounds were too toxic or their effects were not sufficient when tested alone to be selected in both conditions. Combination with 5 nM BTZ has decreased the cell toxicity of 7 compounds and increased the effect on R77C-α-SG rescue of 9 compounds above the threshold of 2 standard deviations allowing their selection in the hits list. The virtual ADMET analysis was used to classify the hit compounds according to parameters predicting the potential success of their clinical development. CDKi and HDACi compounds were in the top 10 of this ranking. Among these drugs identified, we report the particular effect of HDACi with 8 HDACi out of 20 tested as able to rescue α-SG at the membrane level. HDACs are zinc-dependent hydrolases removing acetyl groups from the lysine residues of both histone and non-histone proteins (Allfrey et al., 1964). HDACs are well known as epigenetic modulators regulating chromatin condensation and gene transcription (Verdin and Ott, 2015). In the past decade, their enzymatic effects on non-histones protein has been associated with diverse cellular processes such as gene transcription, DNA damage repair, cell division, signal transduction, protein folding, autophagy and metabolism (Narita et al., 2019). Therapeuticaly, HDACi represent drug candidates with broad therapeutic spectrum and are currently predominantly investigated for the treatment of cancer (Mohammad et al., 2019). In this study, we demonstrated the therapeutical interest of HDACi for misfolded protein diseases.

One of the identified HADCi, givinostat, was described in 2005 as a potent pan-HDACi that exerts activity against all types of HDACs and has been mainly investigated in cancer because of its anti-inflammatory effects (Leoni et al., 2005). Here we report that givinostat effect on α-SG membrane expression was not mediated by *SGCA* gene expression regulation nor by proteasome inhibition suggesting an alternative mechanism. Recent studies have demonstrated that autophagy could be an alternative mechanism of ubiquitinated proteins clearance in a compensatory manner of proteasome inhibition (Kocaturk and Gozuacik, 2018). Because we identified synergistic effect on R77C-α-SG membrane rescue and ubiquitin proteins expression after combination of givinostat and BTZ treaments, suggesting the inhibition of two different signaling pathways, we investigated the effect of givinostat on the autophagy. Here, we demonstrated that givinostat induced the accumulation of autophagic components like LC3B and P62 likely associated to autolysosomes containing undegraded materials. Those abnormal autolysosomes reflected a poorly degradative capacity which was supported by the defective maturation of the V-ATPase V0a1 subunit, subsequently required for proper lysosomes acidification. Together, the data propose an inhibition of the autophagic degradative system at a late stage. Among the HDAC proteins, HDAC6 has been related in many studies to the autophagic pathway. In 2002, HDAC6 was firstly described as a microtubule-associated deacetylase (Hubbert et al., 2002) containing an ubiquitin binding zinc finger (Hook et al., 2002). These properties highlighted the capacity of HDAC6 to bind both polyubiquitinated proteins and dynein motors to transport ubiquitinated proteins through microtubules to aggresomes via tubulin deacetylation (Kawaguchi et al., 2003). HDAC6 was then described to facilitate the autophagosome-lysosome fusion during substrate degradation through the recruitment of a F-actin network (Lee et al., 2010b). Here we demonstrated that givinostat induced HSP90 and tubulin acetylation highlighting its inhibitory effect on HDAC6 and suggesting that the inhibitory effect of givinostat on autophagy leading to the R77C-α-SG protein rescue could implicate the HDAC6 activity. We also demonstrated for the first time an interaction between HSP90 and R77C-α-SG by co-immunoprecipitation. Even if we did not observe a reduced interaction between these two proteins after givinostat treatment, excluding a direct effect of givinostat on the HSP90 chaperone activity, this interaction could explain the identification of 9 HSP90i in the primary hits list of the screening. Here, we demonstrated the effect of givinostat on the autophagic pathway through the maturation of the V-ATPase V0a1 subunit required for lysosomes acidification highlighting a potential new implication of HDAC6 for autophagy.

Clinically, givinostat is currently investigated in a phase III clinical trial for Duchenne’s muscular dystrophy (DMD). The beneficial effects of givinostat on DMD were demonstrated by preclinical studies on DMD mice model (mdx mice) treated with escalating doses of givinostat (ranging from 1 to 10 mg/kg/day) for 3.5 months (Consalvi et al., 2013). This study identified an optimal range of doses between 5 and 10 mg/kg/day for givinostat to induce muscle formation, muscle performance restoration and fibrosis prevention, providing the preclinical basis for a translation of givinostat into clinical studies with DMD patients. Recent preliminary clinical data on twenty ambulant young DMD boys revealed that givinostat treatment leads to a significant increase of muscle tissue in the biospies correlated with a decrease of fibrotic tissue as well as substantial reduction of tissue necrosis and fatty replacement (Bettica et al., 2016). Because similar features are also observed in LGMD, we believe that these clinical observations strengthen the use of this treatment to patients with LGMD. Our results describe here an additional effect of givinostat that would have an interest for patients with missense mutations leading to muscular dystrophies in combination with low doses of BTZ. Further preclinical and toxicological *in vivo* studies are required to evaluate positive or negative impact of these combined treatments. Beyond the identification of this potential treatment for LGMD R3 patients, our study pioneerly explore the concept of shared molecular etiology for rare diseases. While 10% of the world population is affected by one or the other of the 7000 rare diseases, the number of underlying etiologies is much smaller. The emerging strategy consisting to target shared molecular mechanisms is currently revolutionizing drug discovery in rare diseases, making possible to target groups of pathologies in the same time (Brooks et al., 2014; Roessler et al., 2021). Here we describe a complete demonstration of this new paradigm by performing a phenotypical combinatorial drug screening on LGMD R3 disease identifying givinotstat as potential treatment and we demonstrate the positive effect of this drug on LGMD R5 and CF.

## Data availability statement

The original contributions presented in the study are included in the article/Supplementary Material, further inquiries can be directed to the corresponding author.

## Author Contributions

I.R. and X.N. were responsible for the experimental design and project management. L.H. performed the cell culture experiments. L.H. developed the screening assay, carried out screening analysis and realized mechanism of action experiments. N.B. developed the lentivirus and provided the LGMD cell models. G.M and J.T. did the screening. N.M., J.D., M.G., D.P. and N. S. performed and analyzed the ADMET analysis. F.D., S.S. and P.F. provided the CFTR cell models and performed the CFTR analysis. C.D. designed and performed the EM experiments and contributed to edit the manuscript. L.H., C.B., M.B. and E.P. provided technical assistance for cell culture and pharmacological studies. L.H., X.N. and I.R. wrote the manuscript.

## Acknowledgments

The authors thank Marc Peschanski (I-Stem, Corbeil-Essonnes) and Carole Kretz (Institut NeuroMyoGène, Lyon) for helpful discussions, the “Imaging and Cytometry Core Facility” of Genethon for technical support and to Genopole Research, Evry, for the purchase of the equipment and the Platform for Immortalization of Human Cells from the Centre of Research in Myology, Institute of Myology, Paris, for providing immortalized MyoD-inducible human fibroblasts. The French CF association Vaincre la Mucoviscidose is thanked for its financial support. We thank John Riordan (University of North Carolina) and Cystic Fibrosis Foundation for antibodies against CFTR from Antibody Distribution Program. The authors thank Daniela Rotin (University of Toronto) for the gift of HEK293 MSR Grip Tite stably coexpressing eYFP (H148Q/I152L) and F508del-CFTR-3HA and UCSF for the gift of CFBE stably expressing F508del-CFTR.

