## Supplemental Material for "Dual blockade of misfolded alpha-sarcoglycan degradation by bortezomib and givinostat combination"

### Supplementary Figure S1

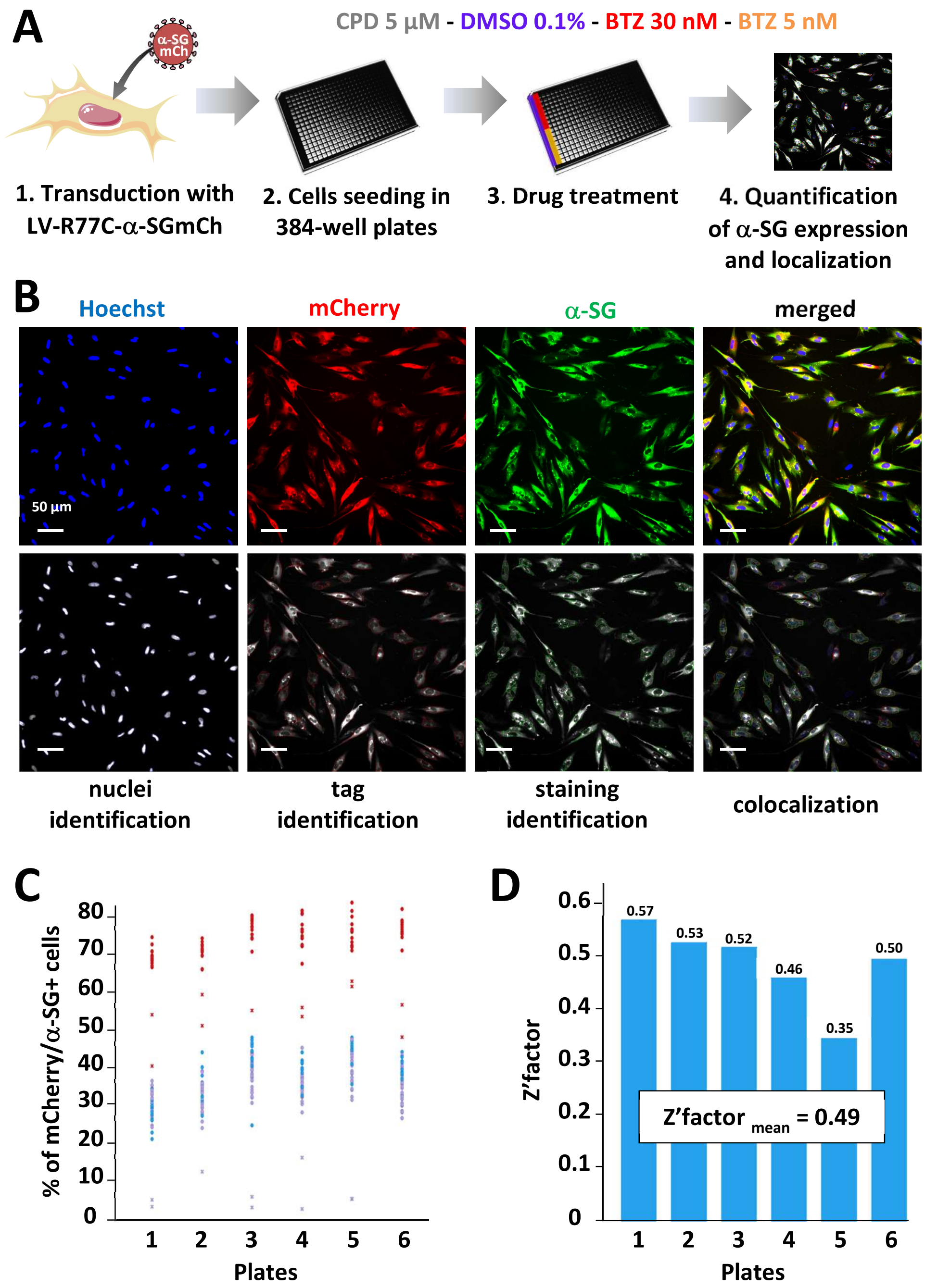


**Supplementary Figure S1. Primary screen cell-based assay for R77C-α-SGmCh membrane rescue (related to Figure 1).** (A) Workflow for the combinatorial high-content screening of R77C-α-SGmCh membrane rescue expression in 384 well plates. (B) mCherry fluorescent signal, membrane α-SG and Hoechst staining in fibroblasts overexpressing R77C-α-SGmCh and treated for 24 hours of treatment with BTZ 30 nM (top panels), and the corresponding masks images (bottom panels) allowing automated quantification of mCherry and membrane α-SG positive cells by colocalization. Scale bar = 50µm. (C) High-throughput screening validation for the R77C-α-SGmCh membrane rescue with the negative control, 0.1% DMSO, and the positive control, 30 nM BTZ, in each of the 384-well plates of the screening. (D) Determination of the Z′ factor in each of the 384-well plates of the screening. CPD = tested compounds; BTZ = bortezomib.

#### Supplementary Figure S2

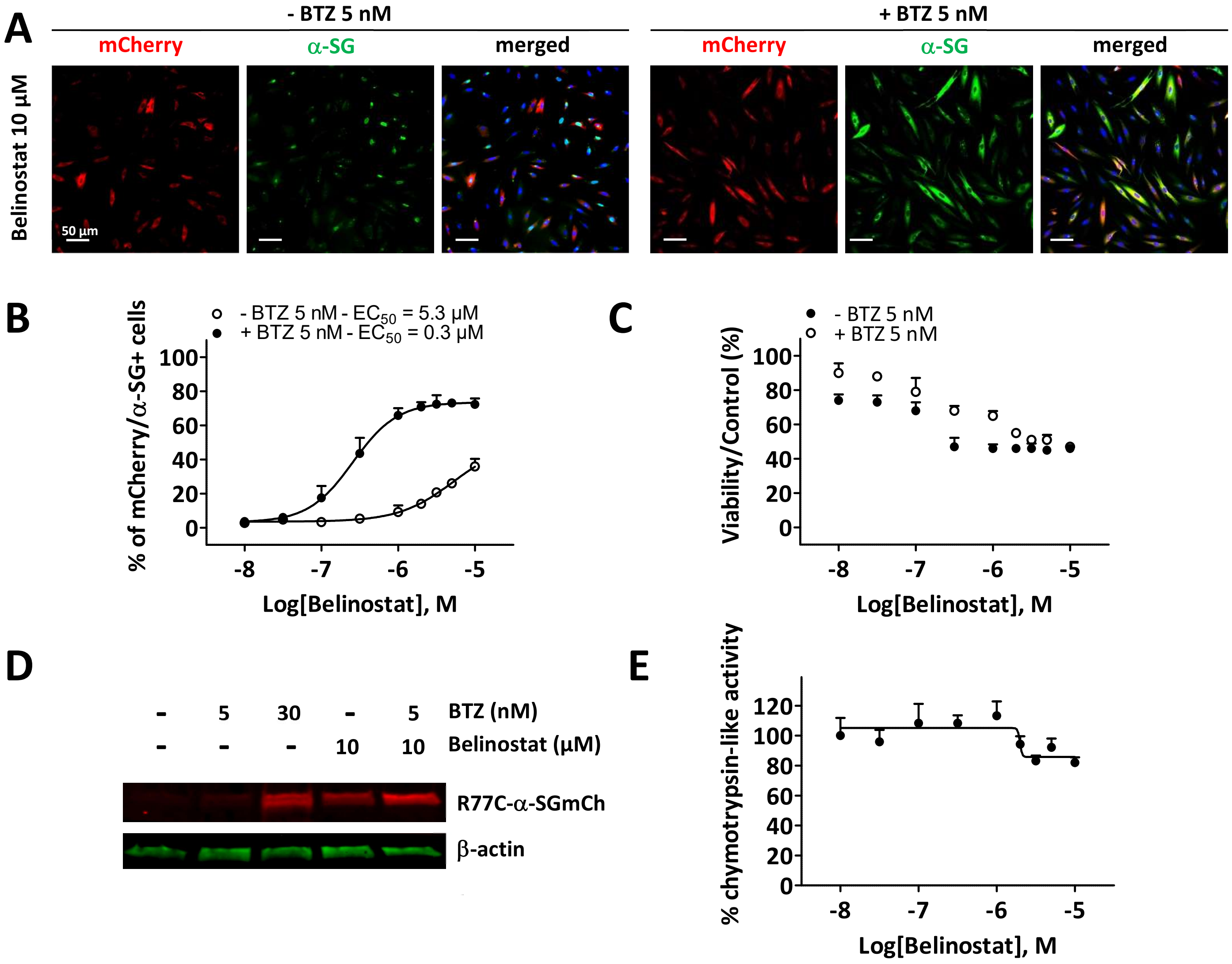


**Supplementary Figure S2. Combinatorial effects of bortezomib and belinostat (related to Figure 2).** Fibroblasts overexpressing R77C-α-SGmCh were treated with DMSO (0.1%), BTZ (5 nM or 30 nM) or belinostat (10 µM or increasing concentrations) alone or in combination with 5 nM BTZ for 24 hours. (A) mCherry fluorescent signal (red) and α-SG staining (green) after fibroblasts treatments. Nuclei are labelled by Hoechst staining (blue). Scale bar = 50 µm. (B, C) Quantification of mCherry and membrane α-SG positive fibroblasts (B) and cell viability (C) following fibroblasts treatments. Each point represents the mean±SD (n=4) of a representative experiment over 3 independent experiments. (D) Immunoblot analysis of α-SG expression after fibroblasts treatments. β-actin was used to evaluate the loading. (E) Quantification of the chymotrypsin-like activity of proteasome following fibroblasts treatments. Each point represents the mean±SD (n=4) of a representative experiment over 3 independent experiments. BTZ = bortezomib.

### Supplementary Figure S3

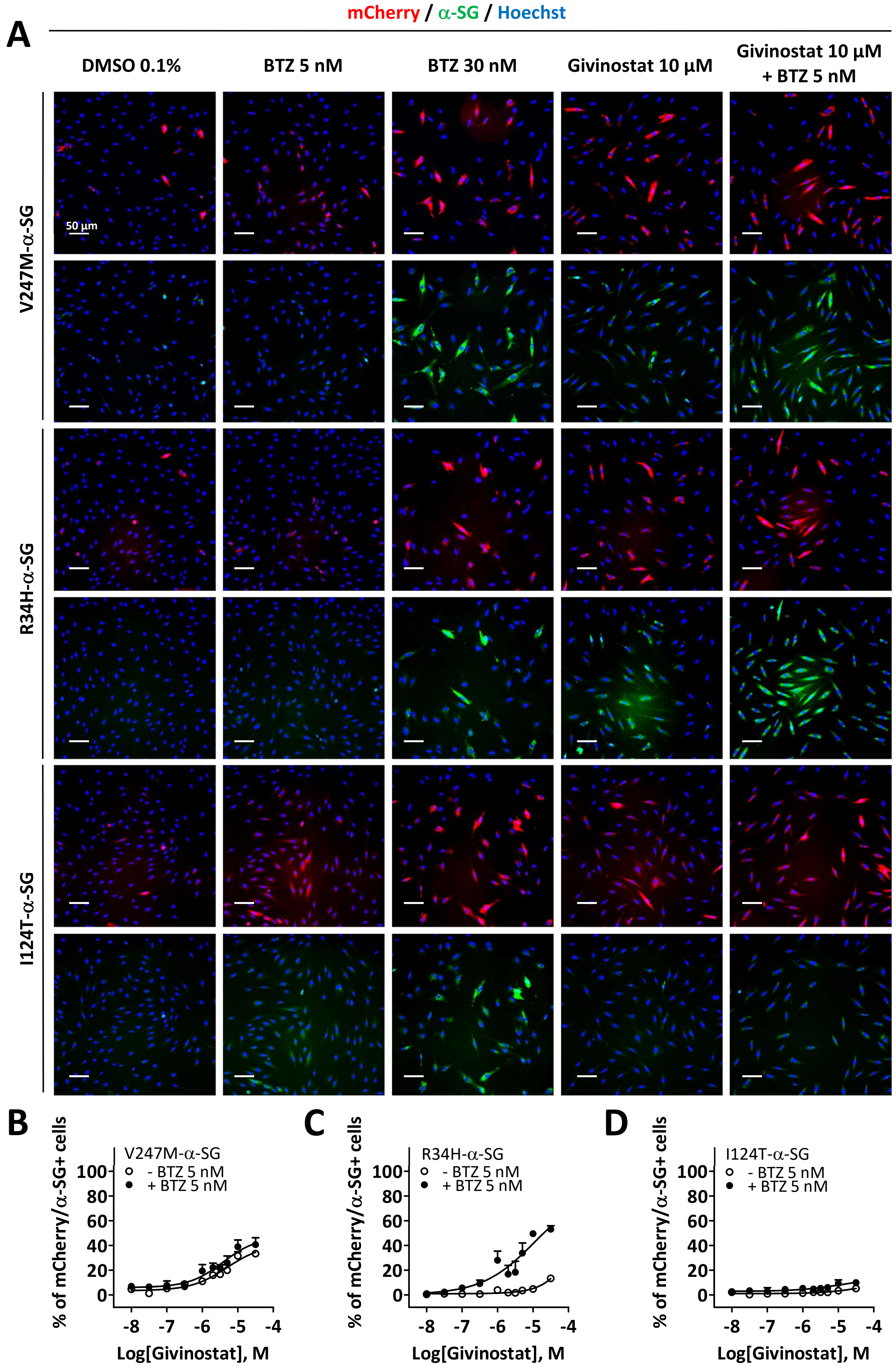

**Supplementary Figure S3 Evaluation of givinostat and bortezomib combination on other missense α-SG mutants (related to Figure 3).** LGMDR3 patient's fibroblasts were transduced with lentivirus expressing R34H-, I124T- or V247M-α-SGmCh constructs and treated for 24 hours with DMSO (0.1 %), BTZ (5 nM and 30 nM), or increasing concentrations of givinostat alone or in combination with 5 nM BTZ. The membrane α-SG expression was monitored by immunofluorescence under non-permeabilized condition. (A) Images of mCherry fluorescent signal (red), membrane α-SG (green) and Hoechst staining (blue) after fibroblasts treatments. Scale bar = 50 µm. (B-D) Quantification of mCherry and membrane R34H-α-SG (B), I124T-α-SG (C) and V247M-α-SG (D) positive cells following fibroblasts treatments. Each point represents the mean±SD (n=4) of a representative experiment over 3 independent experiments. BTZ = bortezomib.

#### Supplementary Figure S4

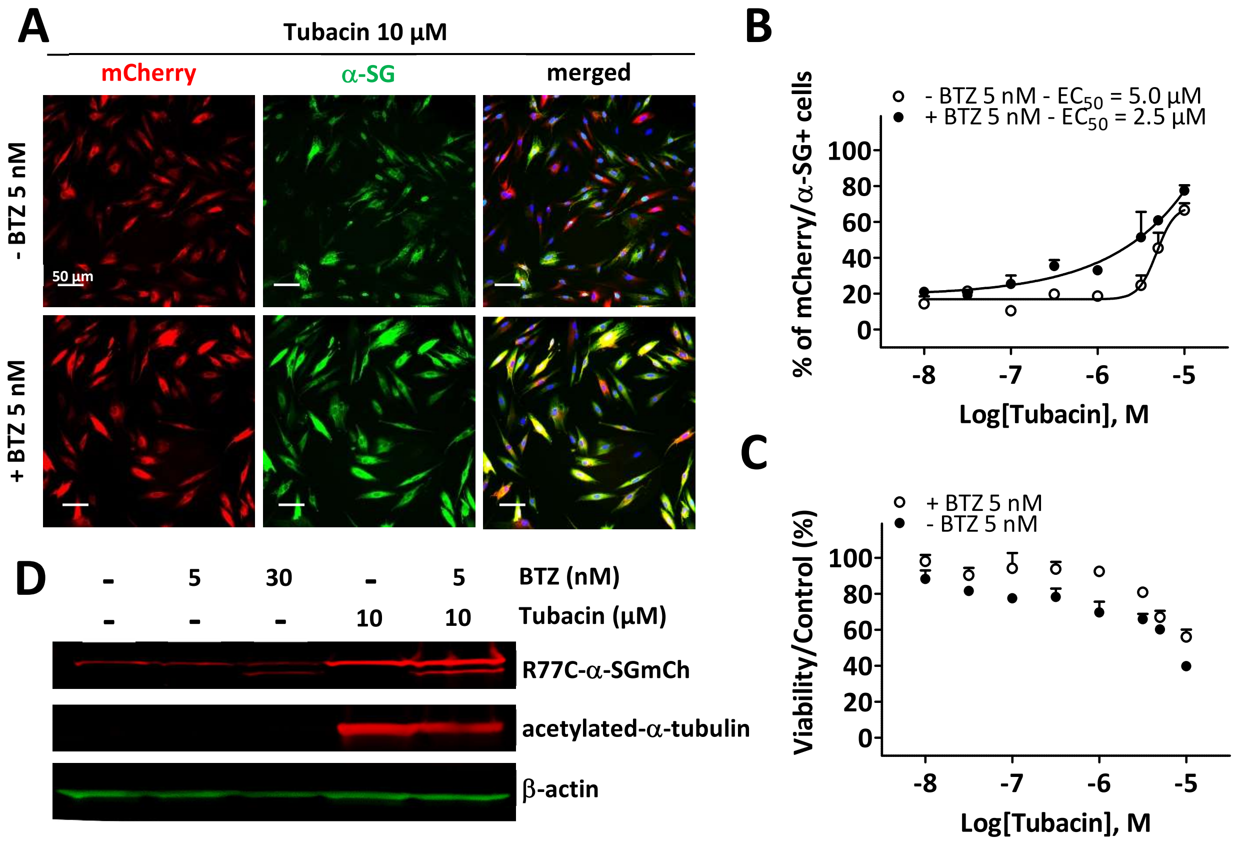


**Supplementary Figure S4. Rescue of misfolded R77C-α-SG degradation by tubacin (related to Figure 5).** (A-D) Fibroblasts overexpressing R77C-α-SGmCh were treated with DMSO (0.1%), BTZ (5 nM or 30 nM) or tubacin (10 µM or increasing concentrations) alone or in combination with 5 nM BTZ for 24 hours. (A) mCherry fluorescent signal (red) and α-SG staining (green) after fibroblasts treatments. Nuclei are labelled by Hoechst staining (blue). Scale bar = 50 µm. (B, C) Quantification of mCherry and membrane α-SG positive fibroblasts (B) and cell viability (C) following treatments. Each point represents the mean±SD (n=4) of a representative experiment over 3 independent experiments. (D) Immunoblot analysis of α-SG and acetylated-α-tubulin expression in fibroblasts after treatments. β-actin was a loading control.

### Supplementary Figure S5

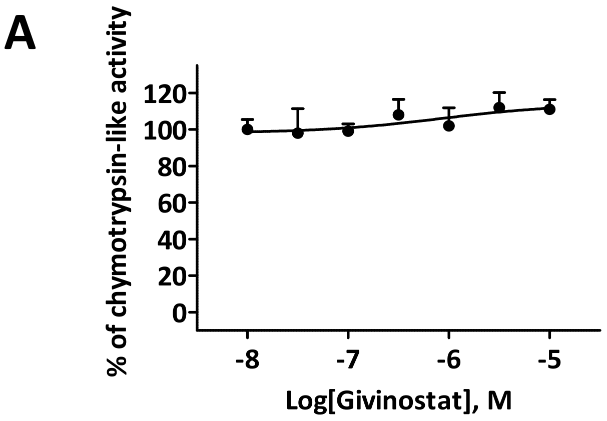


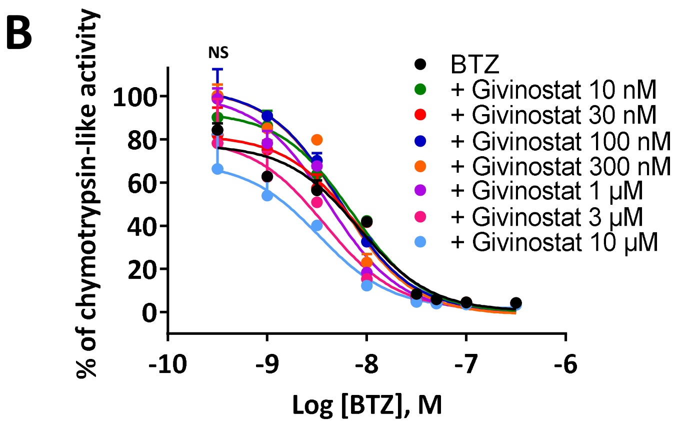


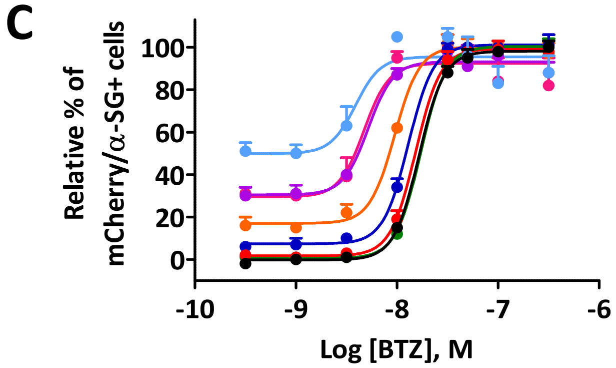


D

| Compound | α-SG $EC_{50}$ (nM) | proteasome $IC_{50}$ (nM) |
|---|---|---|
| BTZ | 16.9 | 8.3 |
| + Givinostat 10 nM | 16.9 | 7.1 |
| + Givinostat 30 nM | 14.8 | 7.9 |
| + Givinostat 100 nM | 12.5 | 5.3 |
| + Givinostat 300 nM | 9.4 | 5.3 |
| + Givinostat 1 µM | 5.2 | 4.1 |
| + Givinostat 3 µM | 3.5 | 3.7 |
| + Givinostat 10 µM | 3.3 | 3.3 |

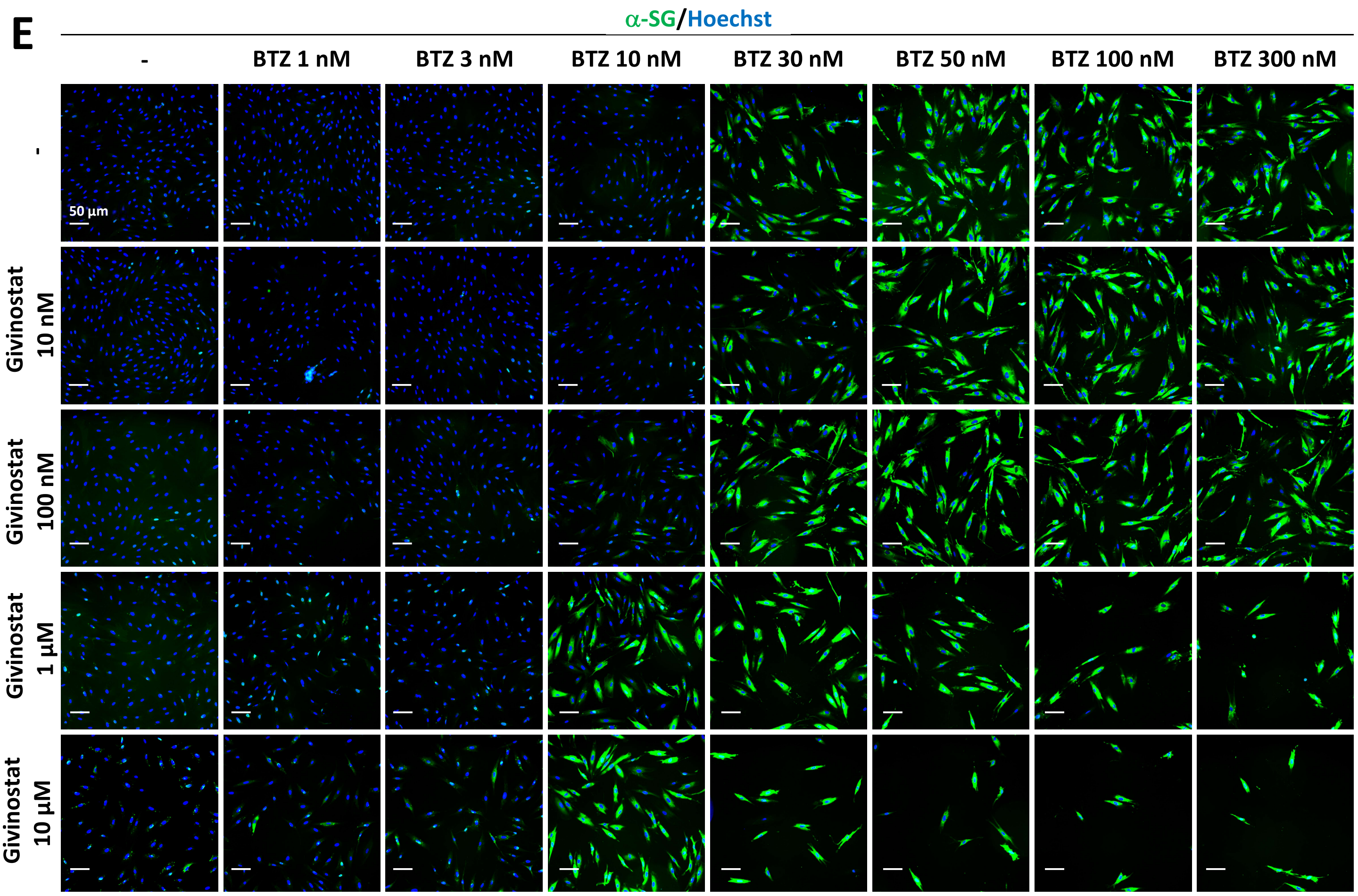

**Supplementary Figure S5. Combination of givinostat and bortezomib effects.** (A-C) Quantification of chymotrypsin-like activity of proteasome (A-B) and mCherry fluorescent signal and membrane α-SG staining (C) in fibroblasts overexpressing R77C-α-SGmCh and treated with increasing concentrations of givinostat alone or in combination with increasing concentration of BTZ for 24 hours.. Values are expressed as percentage of the response induced by 0.1% DMSO (A-B) or as percentage of the maximal response induced by BTZ (C) and each point represents the mean±SD (n=4) of a representative experiment over 3 independent experiments; NS=non-significant (Student's t-test). (D) $EC_{50}$ and $IC_{50}$ of BTZ and combination treatments on membrane α-SG rescue and proteasome activity. (E) Images of membrane α-SG (green) and Hoechst staining (blue) after treatments. Scale bar = 50 µm; BTZ = bortezomib.

#### Supplementary Figure S6

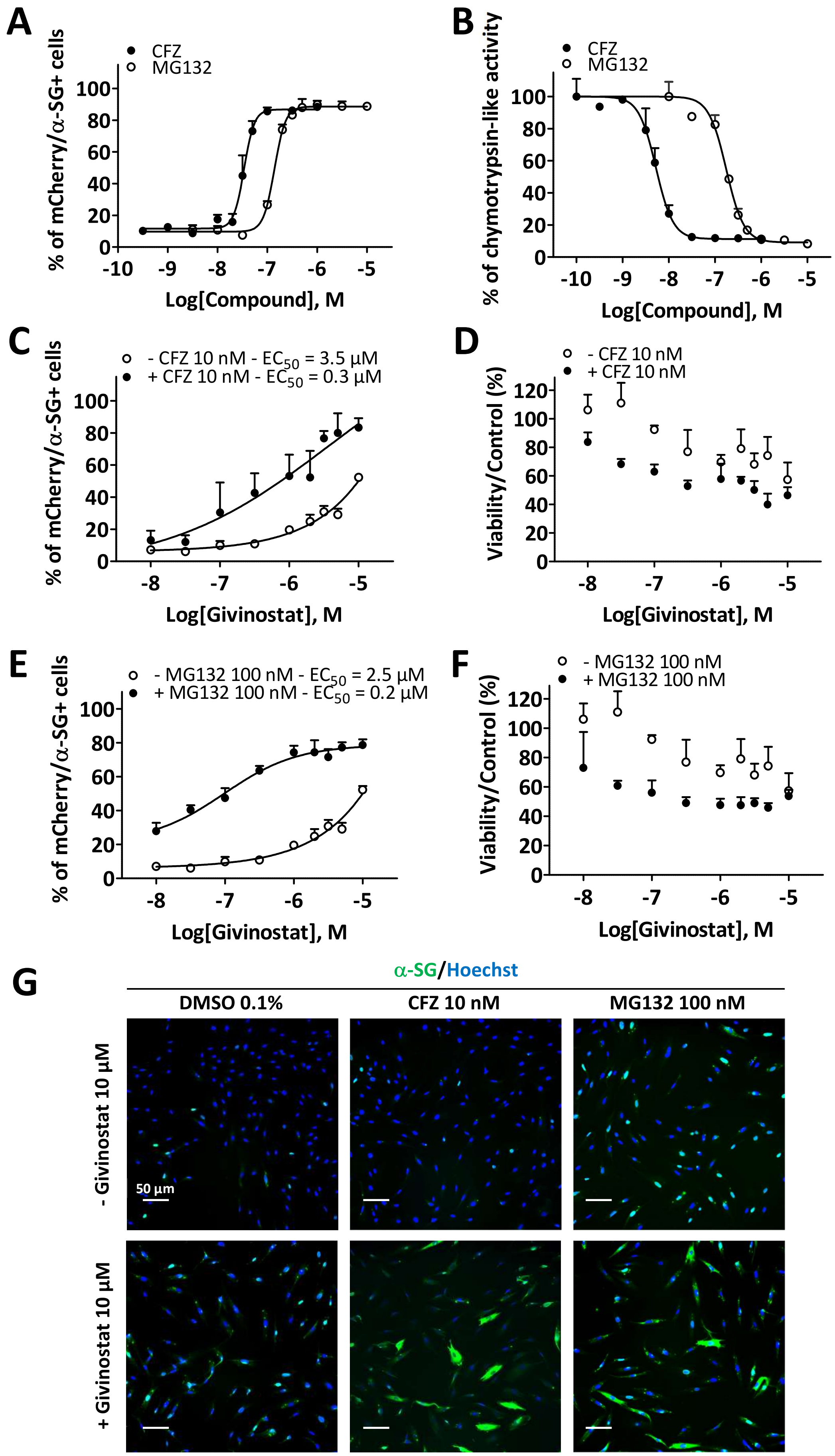

**Supplementary Figure S6. Combinatorial effects of givinostat with the proteasome inhibitors carfilzomib and MG132.** (A-B) Quantification of mCherry and membrane $\alpha$-SG positive fibroblasts (A) or chymotrypsin-like activity of proteasome (B) after fibroblasts treatment with increasing concentrations of CFZ or MG132. (C-G) Quantification of mCherry and membrane $\alpha$-SG positive fibroblasts (C, E), cell viability (D, F) and representative fluorescence images (G) of membrane $\alpha$-SG (green) and Hoechst staining (blue) after fibroblasts treatments with increasing concentrations of givinostat alone or in combination with 10 nM CFZ (C, D, G) or 100 nM MG132 (E, F, G) for 24 hours. Each point represents the mean±SD (n=4) of a representative experiment over 3 independent experiments. BTZ = bortezomib, CFZ = carfilzomib.

### Supplementary Figure S7

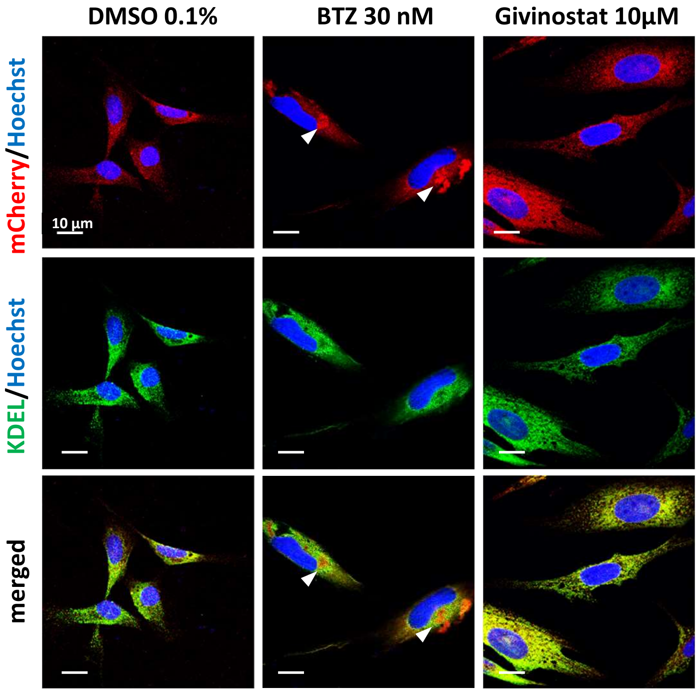


**Supplementary Figure S7. R77C-α-SG aggregation in reticulum endoplasmic under proteasome inhibition (related to Figure 6).** (A) Confocal images of mCherry fluorescent signal (red), KDEL (green) and Hoechst staining (blue) after DMSO (0.1%), BTZ (30 nM) or givinostat (10 µM) treatments. Scale bar = 10 µm.

#### Supplementary Table S1

List of the primers used to introduce the R34H, R77C, I124T, V247M, E263K and C283Y mutations (related to Materials and Methods).

| **Primer name** | **Primer Sequence** |
|---|---|
| R34H Forward | 5'-CACCCACTTGTGGGCCACGTCTTTGTGCACACC-3' |
| R34H Reverse | 5'-GGTGTGCACAAAGACGTGGCCCACAAGTGGGTG-3' |
| R77C Forward | 5'-GCCCCGGTGGCTCTGCTACACCCAGCGC-3' |
| R77C Reverse | 5'-GCGCTGGGTGTAGCAGAGCCACCGGGGC-3' |
| I124T Forward | 5'-CTGGTGCTGGAGACTGGGGACCCAGAA-3' |
| I124T Reverse | 5'-TTCTGGGTCCCCAGTCTCCAGCACCAG-3' |
| V247M Forward | 5'-GTTGACTGGTGCAATATGACCCTGGTGGATA-3' |
| V247M Reverse | 5'-TATCCACCAGGGTCATATTGCACCAGTCAAC-3' |
| E263K Forward | 5'- GCTCACAGAGCCTCTACAAAATCTGTGTGTGTCC-3' |
| E263K Reverse | 5'- GGACACACACAGATTTTGTAGAGGCTCTGTGAGC-3' |
| C283Y Forward | 5'- GGTGTGAGCACCACGTACCAGGAGCACAAC-3' |
| C283Y Reverse | 5'- GGTTGTGCTCCTGGTACGTGGTGCTCACACC-3' |

### Supplementary Table S2

List of the compounds identified as primary hits during the screening in absence or in presence of 5 nM BTZ and the corresponding percentage of R77C-a-SG rescue. Z-Score and percentage of cell viability. Compounds were considered as primary hits when their Z-Score were superior at 2 standard deviations and their percentage of viability superior at 45% (related to Figure 1).
** values beyond selected thresholds*

| | Compound | Family | - BTZ 5 nM | | | + BTZ 5 nM | | |
|---|---|---|---|---|---|---|---|---|
| | | | % rescue | Z-Score | % viability | % rescue | Z-Score | % viability |
| Primary hits only in absence of 5 nM BTZ | Istradefylline (KW-6002) | other | 93.30 | 3.43 | 49.88 | 67.83 | 2.50 | 38.66* |
| | PHA-665752 | other | 88.88 | 3.37 | 57.14 | 81.49 | 2.94 | 26.19* |
| | Triptolide | other | 78.36 | 2.94 | 47.58 | 52.87 | 1.92* | 58.87 |
| | PF 477736 | Kinasei | 76.4 | 2.92 | 48.63 | 73.28 | 2.64 | 41.10* |
| | Benazepril hydrochloride | other | 73.22 | 2.81 | 57.73 | 40.53 | 1.43* | 70.56 |
| | PF-562271 | Kinasei | 62.19 | 2.41 | 57.2 | 39.6 | 1.40* | 55.33 |
| | Dasatinib (BMS-354825) | Kinasei | 60.90 | 2.37 | 93.32 | 41.47 | 1.47* | 79.08 |
| | VU 0364770 | other | 57.49 | 2.26 | 52.59 | 25.04 | 0.83* | 53.14 |
| Primary hits only in presence of 5 nM BTZ | JNJ-26481585 | HDACi | 107.19 | 4.02 | 38.84* | 95.93 | 3.47 | 45.69 |
| | Crizotinib (PF-02341066) | Kinasei | 83.83 | 3.19 | 44.20* | 106.57 | 3.86 | 47.89 |
| | AUY922 (NVP-AUY922) | HSP90i | 81.42 | 3.10 | 38.60* | 73.60 | 2.65 | 48.81 |
| | ENMD-2076 | AKi | 80.19 | 3.06 | 38.80* | 61.17 | 2.19 | 50.59 |
| | AZD7762 | CDKi | 77.76 | 2.97 | 40.12* | 74.97 | 2.70 | 49.88 |
| | BIIB021 | HSP90i | 74.43 | 2.85 | 39.88* | 59.39 | 2.13 | 45.08 |
| | Ganetespib (STA-9090) | HSP90i | 72.40 | 2.78 | 39.53* | 77.37 | 2.79 | 46.46 |
| | PHA-793887 | CDKi | 68.44 | 1.95* | 54.66 | 104.94 | 3.36 | 48.62 |
| | TG101209 | Kinasei | 67.13 | 1.90* | 51.86 | 92.89 | 2.94 | 46.74 |
| | ENMD-2076 L-(+)-Tartaric acid | AKi | 64.54 | 1.81* | 57.11 | 70.53 | 2.15 | 58.57 |
| | Fedratinib ( TG101348) | Kinasei | 62.71 | 1.75* | 73.21 | 98.28 | 3.13 | 50.18 |
| | PF-04929113 (SNX-5422) | HSP90i | 60.23 | 1.66* | 49.07 | 79.49 | 2.46 | 50.44 |
| | Cyclocytidine HCl | other | 50.27 | 1.99* | 49.46 | 60.63 | 2.17 | 45.64 |
| | BX-912 | Kinasei | 50.01 | 1.98* | 60.58 | 61.62 | 2.21 | 56.57 |
| | Foretinib (GSK1363089. XL880) | other | 43.10 | 1.74* | 66.87 | 74.47 | 2.68 | 52.23 |
| | Geldanamycin | HSP90i | 35.37 | 0.78* | 49.31 | 71.33 | 2.17 | 49.97 |
| Primary hits in both conditions | CUDC-907 | HDACi | 103.98 | 3.20 | 50.29 | 105.04 | 3.37 | 46.01 |
| | SB939 (Pracinostat) | HDACi | 99.38 | 3.04 | 59.96 | 88.65 | 2.79 | 56.70 |
| | AZD5438 | CDKi | 94.93 | 2.88 | 48.97 | 88.81 | 2.79 | 56.64 |
| | Dinaciclib (SCH727965) | CDKi | 94.08 | 2.85 | 56.97 | 104.81 | 3.36 | 56.90 |
| | AT9283 | AKi | 90.21 | 3.41 | 48.87 | 96.00 | 3.47 | 51.41 |
| | ITF2357 (Givinostat) | HDACi | 88.24 | 2.65 | 59.17 | 102.93 | 3.29 | 49.87 |
| | JTC-801 | other | 85.63 | 2.56 | 59.42 | 75.51 | 2.32 | 50.18 |
| | AR-42 (HDAC-42) | HDACi | 85.61 | 3.18 | 52.69 | 76.99 | 2.86 | 56.58 |
| | M344 | HDACi | 80.68 | 3.07 | 71.30 | 78.27 | 2.82 | 55.85 |
| | Flavopiridol (Alvocidib) HCl | CDKi | 80.43 | 2.37 | 52.70 | 105.72 | 3.39 | 46.85 |
| | AT7519 | CDKi | 78.83 | 2.32 | 58.29 | 96.22 | 3.05 | 46.33 |
| | Azilsartan Medoxomil (TAK-491) | other | 78.58 | 2.95 | 47.23 | 64.02 | 2.35 | 51.94 |
| | TW-37 | other | 78.35 | 2.99 | 66.58 | 76.65 | 2.76 | 60.81 |
| | 17-DMAG HCl (Alvespimycin) | HSP90i | 77.22 | 2.95 | 52.37 | 79.21 | 2.85 | 46.26 |
| | 17-AAG (Tanespimycin) | HSP90i | 76.60 | 2.93 | 53.05 | 80.15 | 2.89 | 51.51 |
| | SNX-2112 | HSP90i | 73.87 | 2.14 | 55.54 | 77.27 | 2.38 | 53.41 |
| | AT13387 | HSP90i | 73.68 | 2.83 | 51.97 | 57.95 | 2.07 | 49.62 |
| | MLN8237 (Alisertib) | AKi | 72.86 | 2.80 | 57.63 | 68.17 | 2.45 | 47.99 |
| | PCI-24781 | HDACi | 68.61 | 2.65 | 56.59 | 67.72 | 2.43 | 51.56 |
| | Belinostat (PXD101) | HDACi | 67.89 | 2.62 | 58.22 | 62.16 | 2.23 | 60.09 |
| | JNJ-7706621 | CDKi | 64.13 | 2.49 | 56.84 | 71.1 | 2.56 | 52.38 |
| | Danusertib (PHA-739358) | AKi | 59.88 | 2.33 | 63.43 | 75.14 | 2.71 | 55.29 |
| | SNS-314 Mesylate | AKi | 57.26 | 2.24 | 58.95 | 64.37 | 2.31 | 53.20 |

#### Supplementary Table S3

Classification of the hit compounds based on a virtual ADMET analysis using artificial intelligence. Final score, ADMET score, activity score and viability score are detailed. (related to Figure 1).

| Ranking | Compounds | Family | Final score | ADMET score | Activity score | Viability score |
|---|---|---|---|---|---|---|
| 1 | PHA-793887 | CDKi | 0.395 | 0.773 | 1.049 | 0.486 |
| 2 | Dinaciclib (SCH727965) | CDKi | 0.372 | 0.693 | 0.941 | 0.570 |
| 3 | M344 | HDACi | 0.368 | 0.640 | 0.807 | 0.713 |
| 4 | AZD5438 | CDKi | 0.329 | 0.707 | 0.949 | 0.490 |
| 5 | AT7519 | CDKi | 0.325 | 0.533 | 0.788 | 0.583 |
| 6 | SB939 (Pracinostat) | HDACi | 0.318 | 0.587 | 0.994 | 0.600 |
| 7 | CUDC-907 | HDACi | 0.307 | 0.573 | 1.040 | 0.503 |
| 8 | ITF2357 (Givinostat) | HDACi | 0.299 | 0.640 | 0.882 | 0.592 |
| 9 | Istradefylline (KW-6002) | other | 0.298 | 0.547 | 0.933 | 0.499 |
| 10 | TW-37 | other | 0.285 | 0.640 | 0.784 | 0.666 |
| 11 | AT9283 | Aki | 0.282 | 0.760 | 0.902 | 0.489 |
| 12 | JNJ-7706621 | Aki | 0.277 | 0.667 | 0.641 | 0.568 |
| 13 | ENMD-2076 L-(+)-Tartaric acid | Aki | 0.275 | 0.587 | 0.705 | 0.586 |
| 14 | AR-42 (HDAC-42) | HDACi | 0.265 | 0.600 | 0.856 | 0.527 |
| 15 | JNJ-26481585 | HDACi | 0.263 | 0.507 | 0.959 | 0.457 |
| 16 | Crizotinib (PF-02341066) | Kinasei | 0.259 | 0.440 | 1.066 | 0.479 |
| 17 | Dasatinib (BMS-354825) | Kinasei | 0.250 | 0.507 | 0.609 | 0.933 |
| 18 | Fedratinib ( TG101348) | Kinasei | 0.250 | 0.587 | 0.983 | 0.502 |
| 19 | Benazepril hydrochloride | Other | 0.248 | 0.640 | 0.732 | 0.577 |
| 20 | AT13387 | HSP90i | 0.245 | 0.613 | 0.737 | 0.520 |
| 21 | Belinostat (PXD101) | HDACi | 0.242 | 0.600 | 0.679 | 0.582 |
| 22 | PF-04929113 (SNX-5422) | HSP90i | 0.241 | 0.560 | 0.795 | 0.504 |
| 23 | MLN8237 (Alisertib) | Aki | 0.235 | 0.653 | 0.729 | 0.576 |
| 24 | Ganetespib (STA-9090) | HSP90i | 0.235 | 0.573 | 0.774 | 0.465 |
| 25 | 17-AAG (Tanespimycin) | HSP90i | 0.233 | 0.613 | 0.766 | 0.531 |
| 26 | Danusertib (PHA-739358) | Aki | 0.233 | 0.573 | 0.599 | 0.634 |
| 27 | 17-DMAG HCl (Alvespimycin) | HSP90i | 0.232 | 0.533 | 0.772 | 0.524 |
| 28 | TG101209 | Kinasei | 0.232 | 0.453 | 0.929 | 0.467 |
| 29 | JTC-801 | other | 0.231 | 0.453 | 0.856 | 0.594 |
| 30 | PHA-665752 | other | 0230 | 0.613 | 0.889 | 0.571 |
| 31 | AZD7762 | CDKi | 0.229 | 0.587 | 0.750 | 0.499 |
| 32 | PCI-24781 | HDACi | 0.228 | 0.533 | 0.686 | 0.566 |
| 33 | Flavopiridol (Alvocidib) HCl | CDKi | 0.226 | 0.813 | 0.804 | 0.527 |
| 34 | Cyclocytidine HCl | other | 0.225 | 0.547 | 0.606 | 0.456 |
| 35 | SNX-2112 | HSP90i | 0.224 | 0.587 | 0.739 | 0.555 |
| 36 | Triptolide | other | 0.219 | 0.587 | 0.784 | 0.476 |
| 37 | PF 477736 | other | 0.218 | 0.587 | 0.764 | 0.486 |
| 38 | Azilsartan Medoxomil (TAK-491) | other | 0.218 | 0.587 | 0.786 | 0.472 |
| 39 | Geldanamycin | HSP90i | 0.209 | 0.587 | 0.713 | 0.500 |
| 40 | AUY922 (NVP-AUY922) | HSP90i | 0.201 | 0.560 | 0.736 | 0.488 |
| 41 | PF-562271 | Kinasei | 0.190 | 0.533 | 0.622 | 0.572 |
| 42 | VU 0364770 | other | 0.185 | 0.613 | 0.575 | 0.526 |
| 43 | SNS-314 Mesylate | Aki | 0.185 | 0.547 | 0.573 | 0.590 |
| 44 | BIIB021 | HSP90i | 0.182 | 0.680 | 0.594 | 0.451 |
| 45 | ENMD-2076 | AKi | 0.177 | 0.573 | 0.612 | 0.506 |
| 46 | BX-912 | Kinasei | 0.177 | 0.507 | 0.616 | 0.566 |
| 47 | Foretinib (GSK1363089. XL880) | other | 0.171 | 0.440 | 0.745 | 0.522 |
